# Revealing taxonomic signals in plant volatiles with phytochemistry, machine learning and trait mapping

**DOI:** 10.64898/2026.07.28.740953

**Authors:** Anupama Sekhar, Arpana Alka, Ambedkar Dukkipati, Vinita Gowda

## Abstract

Plant volatiles have long been used as taxonomic characters, especially in chemotaxonomy. Exploring the utility of chemotaxonomy has been widely regarded critical for drug discovery, despite the knowledge that phytochemicals are often evolutionarily labile. Despite this, the use of chemotaxonomy has been restricted primarily due to the complex interpretations behind translating chemical characters into phylogenetically operational characters. In this paper, we propose a method that integrates phytochemistry, machine learning, and trait mapping to examine whether plant volatiles retain signals for classification across broad taxonomic ranks and which components of the volatile metabolites carry these signals. Leveraging a global dataset comprising 2,139 volatiles across 429 plant species, we trained classifiers on presence-absence data and molecular fingerprints that can predict species to their taxonomic ranks. Incorporating structural features improves performance, suggesting that plant volatiles may encode lineage-specific chemical signatures. Rather than single diagnostic markers, combinations of volatiles informed taxonomic predictions, indicating biosynthetic constraints within volatile clusters. Fingerprint-derived clusters showed a lineage-dependent pattern when mapped onto phylogeny. The proposed method provides an integrated approach to revisit chemotaxonomy and trait evolution to understand the chemical diversity across lineages. This approach can also support comparative chemical prediction, especially for understudied closely related plant groups.

## INTRODUCTION

Plants synthesise and emit a wide array of structurally diverse volatiles that are highly valued in flavour, fragrance, and pharmaceutical industries. These volatiles may be present in plants due to shared ancestry^[1]^, ecological pressures^[2]^, and environmental factors^[3–4]^. When shared through ancestry, plant volatiles play an important role in chemotaxonomy, in which plants are classified based on their unique chemical signatures^[5–8]^. For example, plant families such as gingers (Zingiberaceae), mint (Lamiaceae), and citrus (Rutaceae) are universally recognisable from their signature volatiles, and the glandular structures responsible for synthesising and storing these volatiles are also similarly conserved within these families.

The distribution of plant volatiles along lineages can arise from diverse evolutionary processes (Fig. 1). For instance, genetic constraints can act on key biosynthetic enzymes and their regulation, resulting in particular pathways or structural features in the most recent common ancestor. When these traits are inherited by the descendants, closely related plant species tend to emit similar volatile profiles, a pattern known as phylogenetic clustering. This has been observed in the synthesis of isoflavones, a subset of phenolics, restricted to the legume subfamily Papilionoideae^[9]^, due to lineage-specific origin of isoflavone synthase^[10–11]^. In contrast, similar volatile profiles in unrelated lineages may be due to convergence under similar ecological or environmental conditions, as observed in the production of cyanogenic glucosides in many plant families as a defense against herbivores and predators^[12]^. The third pattern in plant volatiles is when close relatives show divergence in chemical signatures under variable ecological or environmental conditions, as observed in the genus *Bursera*, where species accumulate greater chemical diversity and complexity in defensive volatiles over evolutionary time, potentially in response to plant-herbivore interactions^[13]^.

**Fig. 1.**
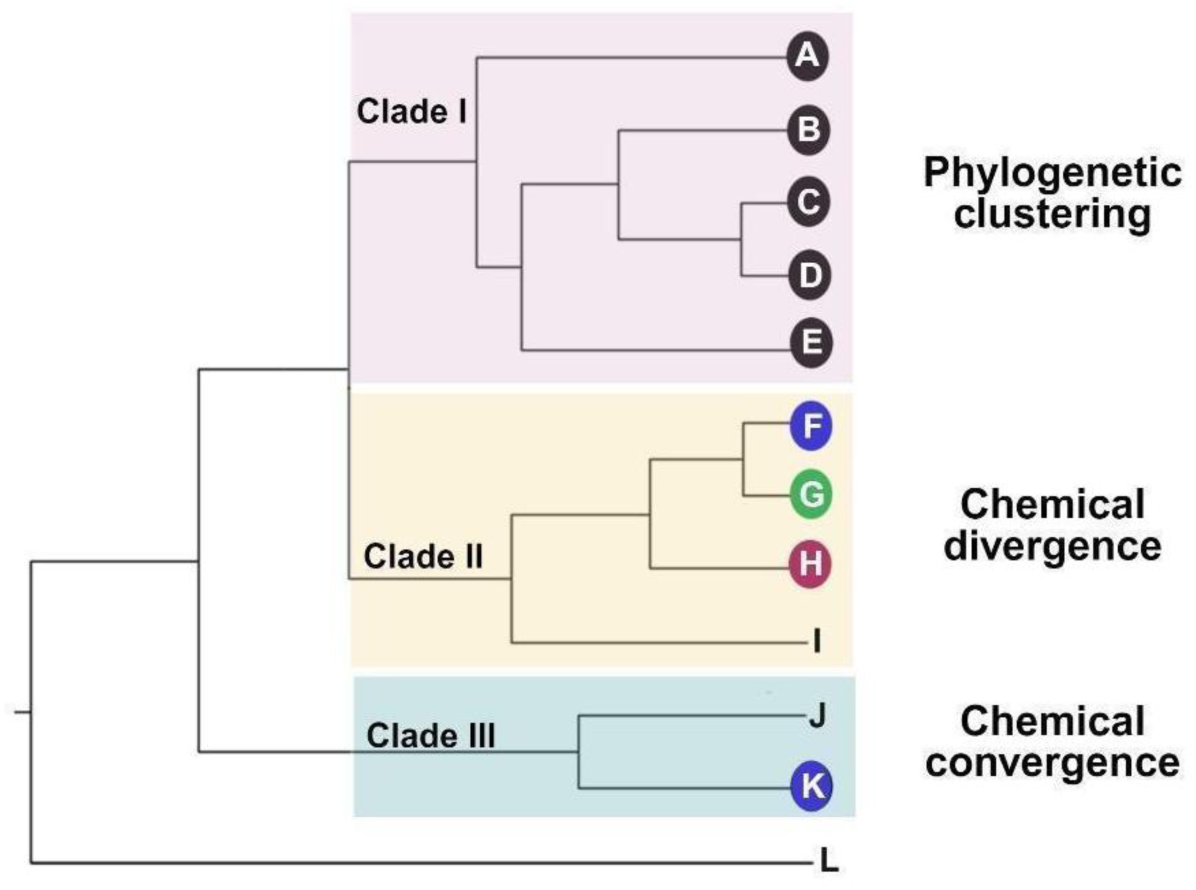
Illustration of the evolutionary patterns of characters (plant volatiles) on a hypothetical phylogeny. A-L represent species on a phylogenetic tree, and circles with the same colour on the tip denote the presence of similar chemicals. All members of Clade I (species A-E) show phylogenetic clustering due to their shared chemistry. In contrast, members of Clade II (F-I) emit different volatiles, indicating divergent evolution. Species F and K from two distant clades emit the same volatiles, showing convergent evolution.

While phylogenetic clustering, convergence, and divergence in chemical data can capture genetic relatedness as well as ecological and environmental adaptations, chemotaxonomic studies have predominantly focused on narrow taxonomic scales ^[14, 15, 16, 17]^, such as within genera or families. Some large-scale studies have examined shared volatile patterns across families^[18]^, and evolutionary signals have been tested for plant-specialized metabolites^[19]^ or specific classes of plant volatiles^[20]^. Since most studies explore utility of chemotaxonomy within a predefined taxonomic rank, that is genus or family, it remains unclear if: a) ranks are defined by diagnostic rare volatiles or by multivariate combinations of volatiles, and b) volatiles retain signals across higher taxonomic ranks.

Plant volatiles can be utilised in taxonomic or phylogenetic studies if they can be treated as morphological characters whose homology is known. However, although volatiles and their classes are known to have distinct structural or functional scaffolds and motifs, their utility has been limited because elucidating their features is very complex. This has resulted in chemotaxonomic studies often relying on spectral features or the presence of volatile classes^[21–22]^ as characters, thus severely underutilising the traits available from plant volatiles.

In the past two decades, structural features of metabolites have been encoded in unique substructural patterns, identified as molecular fingerprints^[23–24]^, which can transform the utility of plant volatiles in evolutionary studies. The fingerprints can be represented as binary bit strings, making them useful for comparative studies. Each bit records the presence or absence of a specific substructure within a metabolite. For instance, in PubChem fingerprints, bits 0 to 114 represent element counts, and bits 115 to 262 encode ring structures, while later bits test for the presence of bonded-atom pairs, bond types, and aromaticity among nearest neighbours in a chemical structure. The combination of the presence of these bits captures the overall chemical structure of a metabolite (Supplementary Fig. S1). Further, recent studies have shown that structurally similar metabolites are typically part of the same biosynthetic pathways^[25–26]^ and may be genetically constrained within plant lineages. For example, betalains, a structurally coherent class of pigments, are produced exclusively in Caryophyllales^[27]^. While the incorporation of molecular fingerprints alongside metabolite-level data is attractive, the structural interpretation of volatile profiles remains a highly complex, high-dimensional problem that is also computationally intensive. Further, since different fingerprints capture variable aspects of the chemical structure^[28–29]^, choosing the appropriate molecular fingerprint is critical for a robust prediction model.

Machine learning (ML) algorithms offer powerful data-driven approaches for analysing high-dimensional data and can uncover subtle patterns. While ML is widely used in the virtual screening of active compounds in drug discovery^[30]^, its application in chemotaxonomy is limited. Recently, unsupervised learning methods have been used to show congruence between genetic relatedness and chemical similarity in plants using one type of fingerprint^[25–26]^. Further, supervised ML using different types of fingerprints has also been shown to be useful within the plant family Ranunculaceae by Chen et al. ^[15]^, although supervised ML approaches that directly train classifiers on plant volatile data are rare.

In the present study, we employed machine learning approaches to identify chemical similarities among plant volatiles and integrated the results with phylogenetic methods to explore the utility of volatiles in distinguishing species to their respective taxonomic ranks. We first tested whether volatiles and their molecular fingerprints can serve as accurate species identifiers in a global qualitative dataset comprising 2,139 plant volatiles from 429 plant species. Next, we evaluated whether the volatile clusters derived from unsupervised learning exhibit phylogenetic structure. Results from our study contributes to providing a structural and computational perspective on chemotaxonomy and trait evolution by treating metabolite structural features as evolutionary traits. When chemical similarity is often used as a proxy for bioactive property similarity^[31]^, this integration can also be relevant to natural product and drug-discovery research. By quantifying how volatile metabolites capture taxonomic information and align with evolutionary relationships, our study has implications for comparative chemical prediction, especially for understudied closely related plant groups.

## RESULTS

### Generation of fingerprint clusters and alignment with volatile classes

Simplified Molecular Input Line Entry System (SMILES) notations were available for 1,668 volatiles, from which 2-10 volatile clusters were generated per fingerprint type using k-means, agglomerative, Density-Based Spatial Clustering of Applications with Noise (DBSCAN), and spectral clustering. The Calinski-Harabasz scores were highest for the SubCount fingerprint type, ranging from 1000 to 1300 (Supplementary Table S2). Next, we assessed how well each fingerprint cluster reflects structural or biosynthetic similarity among volatiles. The highest normalised mutual information (NMI) scores for structural classes (0.508; Supplementary Fig. S2a) were also observed for SubCount type after agglomerative clustering with k=7. The SubCount fingerprint distinguished aromatic and non-aromatic nitrogen and sulphur volatiles as well as sesqui- and diterpenes from mono- and irregular terpenes into distinct clusters (Fig. 2a; Supplementary Fig. S3). When biosynthetic classes were considered, PubChem fingerprints after k-means clustering with k = 3 yielded the highest NMI (0.461) and adjusted rand index (ARI; 0.448) scores (Supplementary Fig. S5). Shikimate-derived aromatics (99.5%), fatty acid derivatives (100%), and terpenoids (72%) formed three distinct clusters derived from PubChem fingerprints (Fig. 2b; Supplementary Fig. S6). Although Molecular ACCess System (MACCS) fingerprint with k=3 after k-means clustering achieved a higher ARI score (0.475) (Supplementary Fig. S5b), the clusters were compositionally mixed, indicating poor interpretability for most of the volatile classes, except for terpenoids (78.6%) (Supplementary Fig. S6). Since NMI scores captured the overall similarity patterns between ML-derived clusters and ground truth labels, for subsequent evolutionary analyses, we used the optimal clusters generated from the fingerprint types that gave the highest NMI scores.

**Fig. 2.**
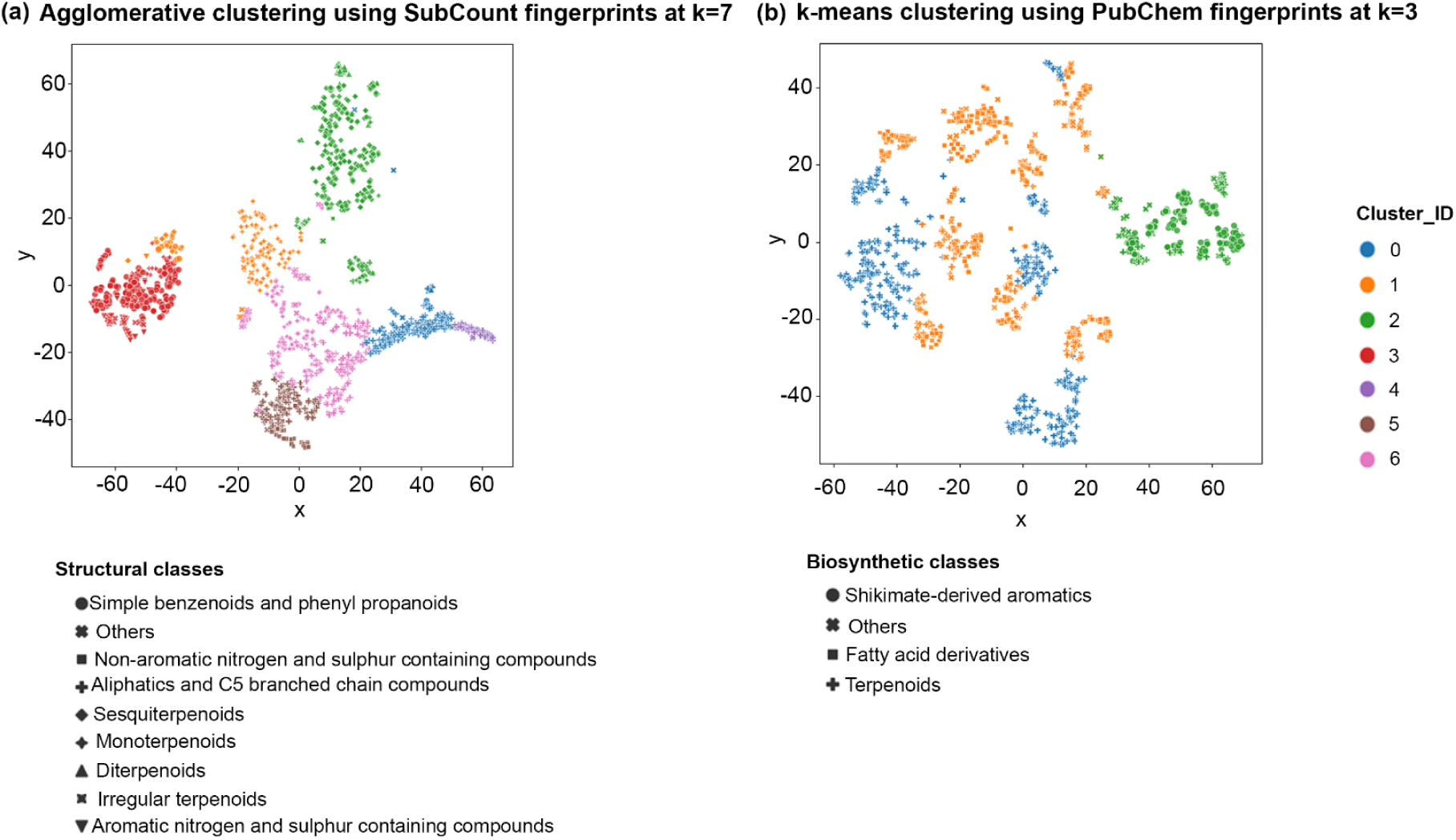
t-distributed stochastic neighbour embedding (t-SNE) plots that show the volatile clusters derived from two fingerprint types. The fingerprints that best reflect the **(a)** “structural” classes using agglomerative clustering of SubCount fingerprints and **(b)** “biosynthetic” classes using k-means clustering of PubChem fingerprints based on the highest NMI scores. Colours represent the cluster assignment, and shapes represent the classes.

### Taxonomic rank predictions using plant volatiles

Table 1 shows that the inclusion of molecular fingerprints in the ML models, along with binary presence-absence of volatiles as features, improved the classification performance for all the taxonomic ranks. The highest classification accuracy was achieved with extended (EXT) fingerprints for order- and family-level predictions with the MultiLayer Perceptron (MLP) model, while Sub fingerprints performed best at the genus level with the Logistic Regression (LR) model.

**Table 1.**
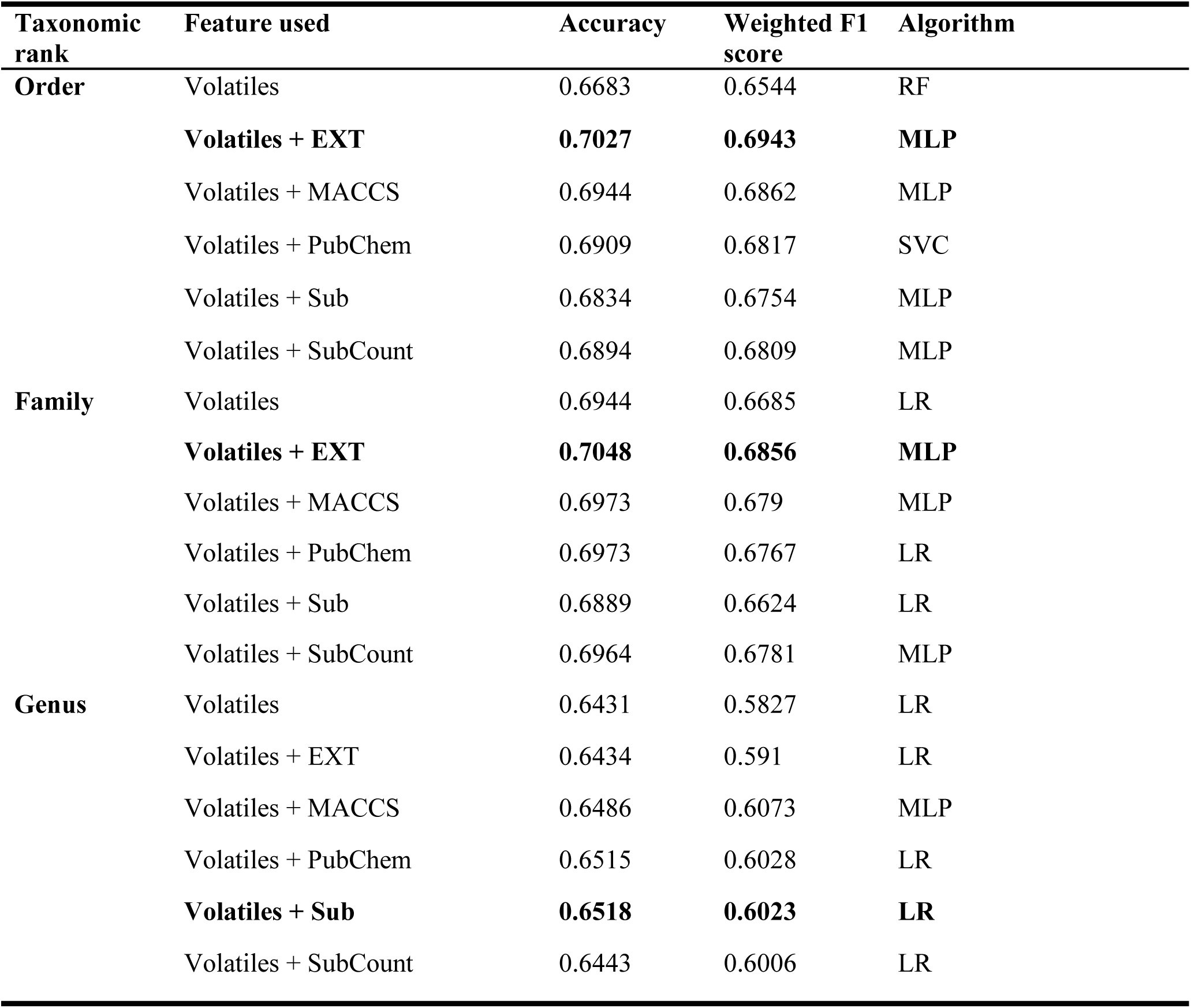
Classification models across taxonomic ranks. Summary of the optimal models obtained after grid search and cross-validation, trained using the presence and absence of plant volatiles and five fingerprint types as features. The model that showed the highest accuracy for each of the taxonomic ranks is represented in bold. Abbreviations: LR - Logistic regression, MLP – Multilayer Perceptron, RF - Random Forest, SVC - Support Vector Classifier.

The classifiers achieved more than 50% recall for 26 plant orders and 40 families (Supplementary Fig. S8), with plant orders with good taxonomic representation (Gentianales, Ranunculales, Caryophyllales, Myrtales, Sapindales, Rosales, Malpighiales, Magnoliales, Asperagales, and Lamiales) also occupying the diagonals in the confusion matrix (Supplementary Note S3; Supplementary Figs. S9, S10). Recall values represent the number of times a plant species is correctly predicted to its taxonomic rank, and lower recall values did not always relate to lower sample support (such as in the cases of Acorales, Liliales, Proteales, Dipsacales, and Vitales; Supplementary Fig. S8a). This pattern indicates that the model accurately predicted plant species to the respective taxonomic ranks, suggesting distinctive chemical profiles. Conversely, 9 orders and 24 families showed poor predictive performance, with a recall below 30% (Supplementary Fig. S8), indicating that the model struggled to learn discriminative patterns for those plant orders and families (Supplementary Note S3; Supplementary Figs. S9, S10).

Additionally, the ML models accurately classified plant species using combinations of volatiles rather than unique, species-specific diagnostic volatiles (Supplementary Notes S4, S5; Supplementary Figs. S12-S15).

### Distribution of fingerprint-derived volatile clusters on a phylogenetic tree

The reconstructed tree topology from Internal Transcribed Spacer (ITS) sequences broadly recovered the expected angiosperm relationships. An exception was Ericales, which belongs to Asterids and forms a separate clade, likely due to limited sampling in that order. To determine the distribution of the volatile clusters, we mapped the fingerprint-derived volatile clusters onto the phylogenetic tree and calculated the D-statistics. We focused on EXT fingerprints, which yielded the best classification performance across taxonomic ranks (Table 1). We selected the optimal number of clusters based on the highest NMI scores for EXT fingerprint type, which showed k=6 after agglomerative clustering for structural classes (Supplementary Fig. S2a) and k=3 after k-means clustering for biosynthetic classes (Supplementary Fig. S5a). The structural groupings comprised two terpene-dominated clusters, one dominated by simple benzenoid, phenylpropanoid, and aromatic nitrogen- and sulphur-containing volatiles, and three clusters enriched in aliphatic and C5 branched chain volatiles. The biosynthetic pathway clusters represent three major volatile origins: volatiles derived from the shikimate pathway, terpenoid pathways, and the lipoxygenase pathway. The EXT fingerprint-derived volatile clusters not only recovered known biosynthetic classes but also revealed finer structural distinctions, including multiple aliphatic clusters (PM3a-c in Fig. 3). We also mapped the distribution of volatile clusters derived from SubCount and PubChem fingerprints that showed the best alignment with the structural and biosynthetic volatile classes, respectively, from the clustering analyses (see Supplementary Table S3 and Supplementary Figs. S16-S17).

**Fig. 3.**
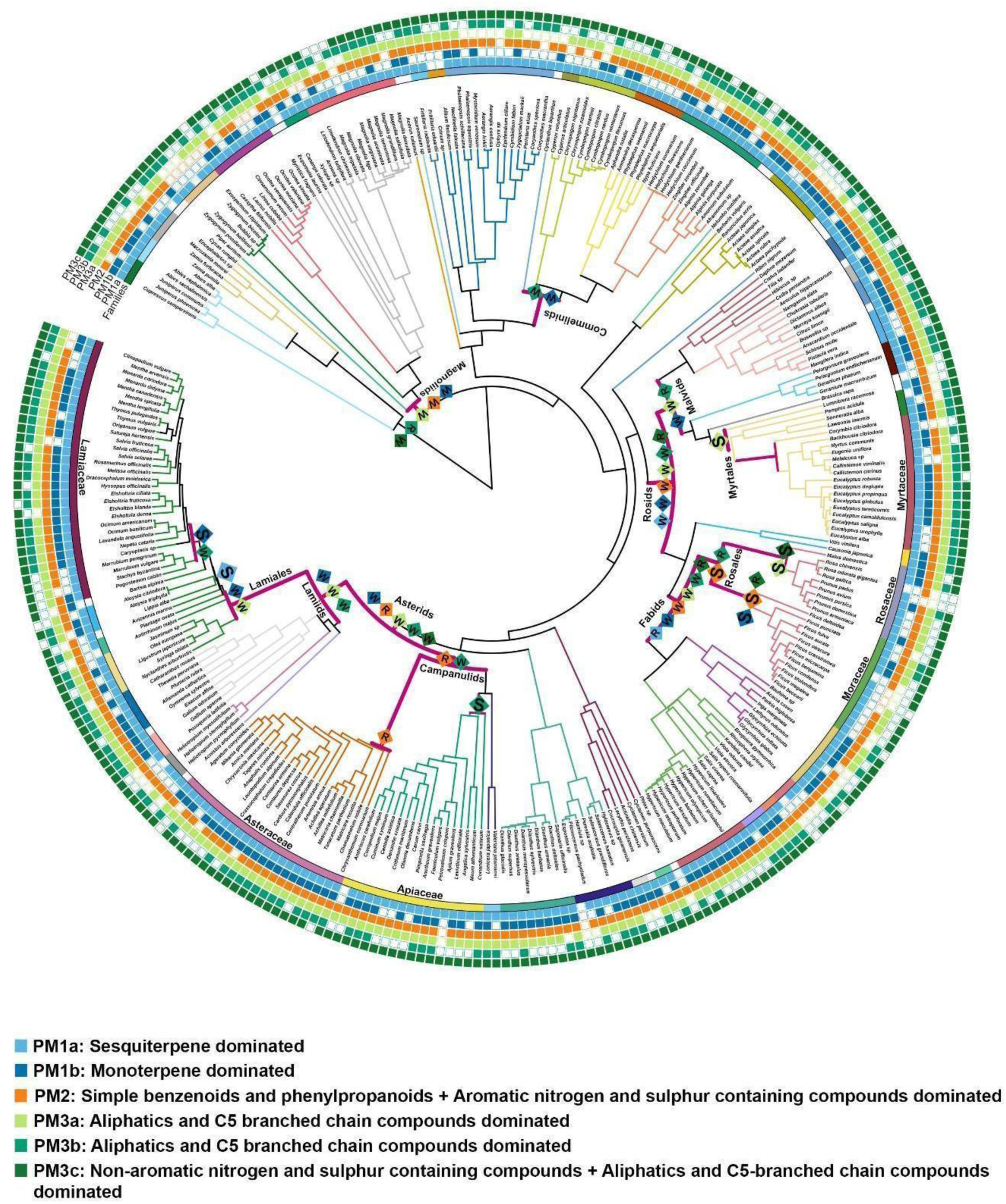
Mapping of EXT fingerprint-derived clusters based on NMI scores of structural classes onto the phylogeny. Coloured branches represent the different orders, and coloured stripes represent different families. Clusters are represented as coloured squares, where presence is indicated by filled squares and absence by unfilled squares. The clades analysed for D-statistics are marked as bold magenta lines. The evolutionary patterns of volatile clusters that gave significant phylogenetic signal for each clade are represented as S = strong phylogenetic clustering, W = weak clustering, R = random shuffle.

Table 2 shows the D-statistics of the volatile clusters that represented structural and biosynthetic classes, generated from the EXT fingerprint, for the entire phylogeny and across specific angiosperm clades. A weak phylogenetic clustering (0 < D < 1) was observed for the entire phylogeny. All the EXT-derived clusters exhibited a distribution that is significantly different from one expected under the Brownian motion model (*p*_Brownian motion_ < 0.05), while all except one cluster showed a distribution significantly different from a random distribution (*p*_random_ < 0.05), with the fatty acid derivative-dominated cluster being the exception. When 11 subsets of angiosperms were considered, most clusters showed weak phylogenetic clustering, random distribution, or no statistically significant phylogenetic signal. Nevertheless, some of the volatile clusters showed a strong phylogenetic clustering (D < 0) in a lineage-specific manner. This included aliphatic and C5 branched chain volatiles dominated clusters in Myrtales, simple benzenoid and phenylpropanoids and aromatic nitrogen and sulphur-containing volatiles dominated clusters in Rosales and sesquiterpene-dominated clusters in Lamiales, among structural classes. The monoterpene-dominated cluster showed a strong signal in Lamiaceae and Moraceae, whereas one of the aliphatic-dominated clusters showed a strong signal in Rosaceae, which either exhibited weak or non-significant signals at corresponding order levels (Fig. 3). Similarly, among biosynthetic classes, a strong phylogenetic clustering was detected in the cluster dominated by shikimate-derived aromatics in Rosales and the terpenoid-dominated cluster in Malvids, although the changes in signal strength across taxonomic ranks were less pronounced than those observed for structural clusters (Fig. 4).

**Fig. 4.**
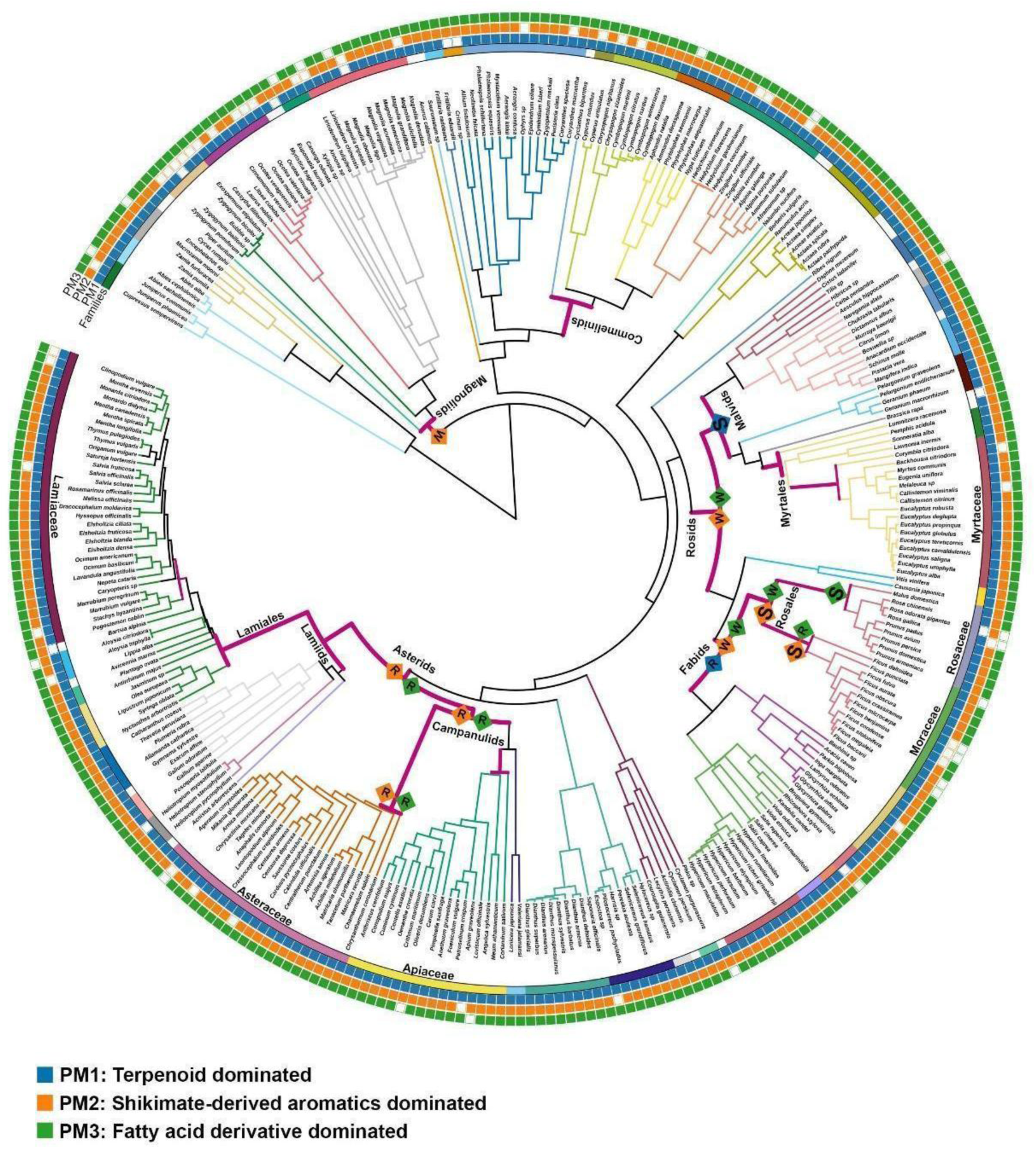
Mapping of EXT fingerprint-derived clusters based on NMI scores of biosynthetic classes onto the phylogeny. Coloured branches represent the different orders, and coloured stripes represent different families. Clusters are represented as coloured squares, where presence is indicated by filled squares and absence by unfilled squares. The clades analysed for D-statistics are marked as bold magenta lines. The evolutionary patterns of volatile clusters that gave significant phylogenetic signal for each clade are represented as S = strong phylogenetic clustering, W = weak clustering, R = random shuffle.

**Table 2.**
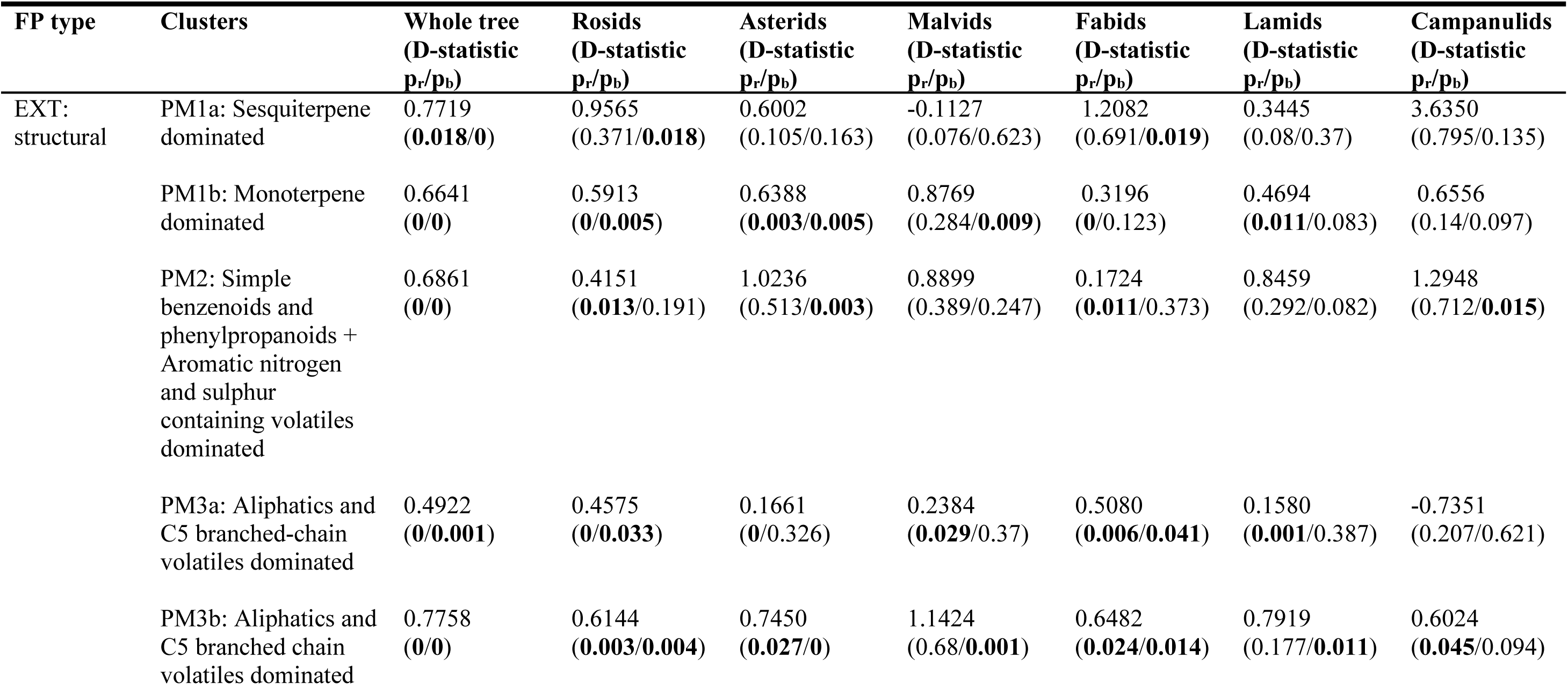
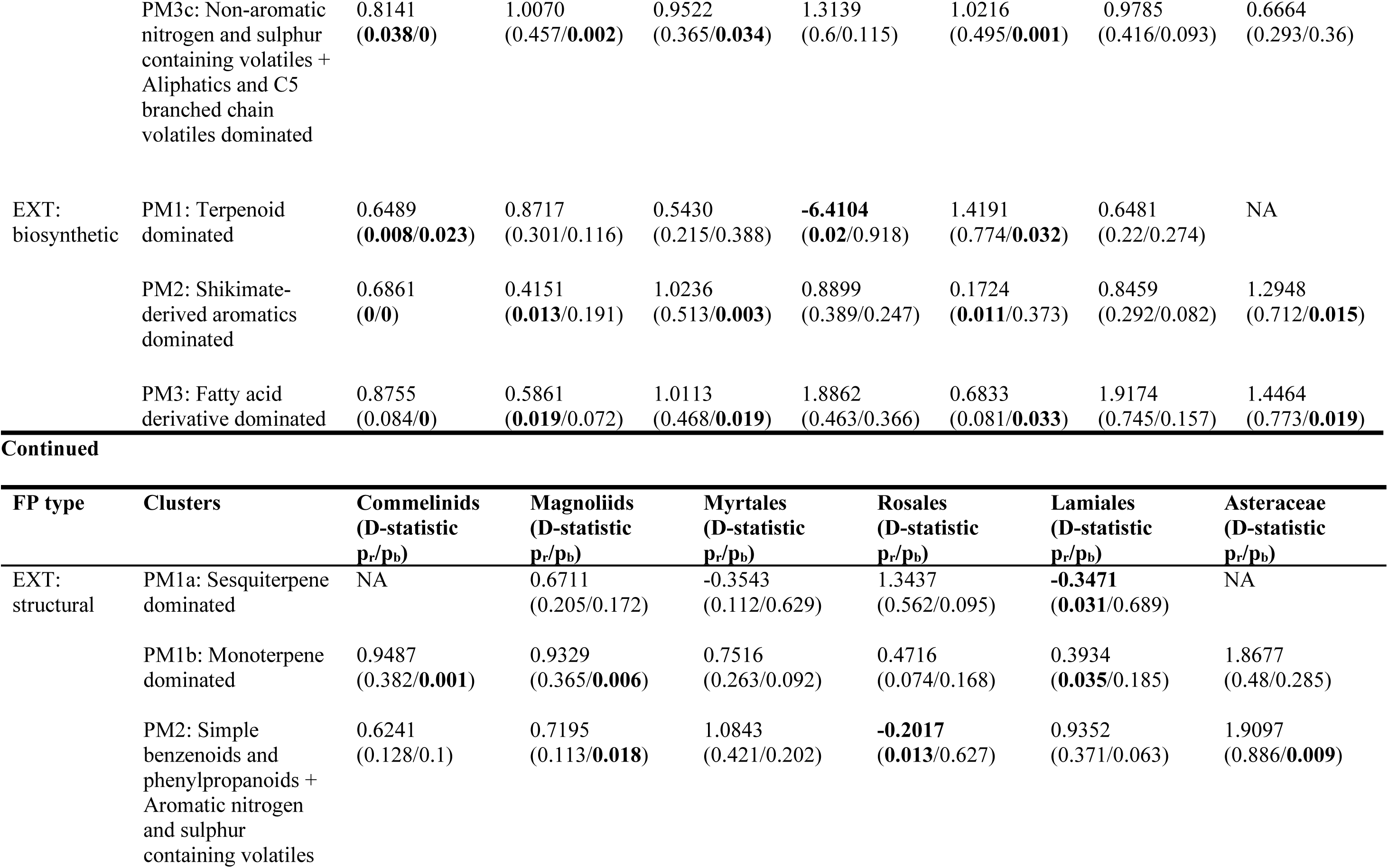
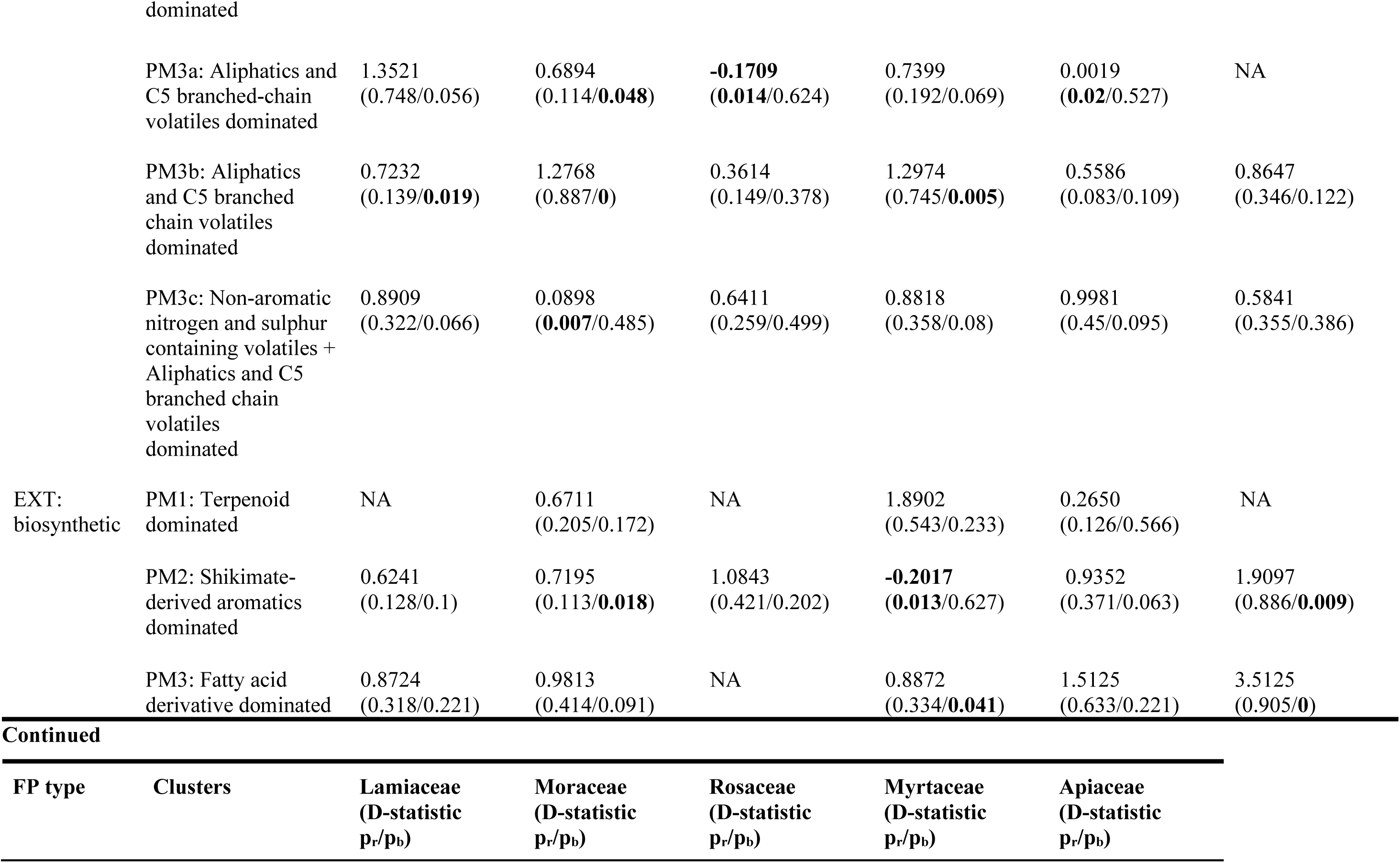
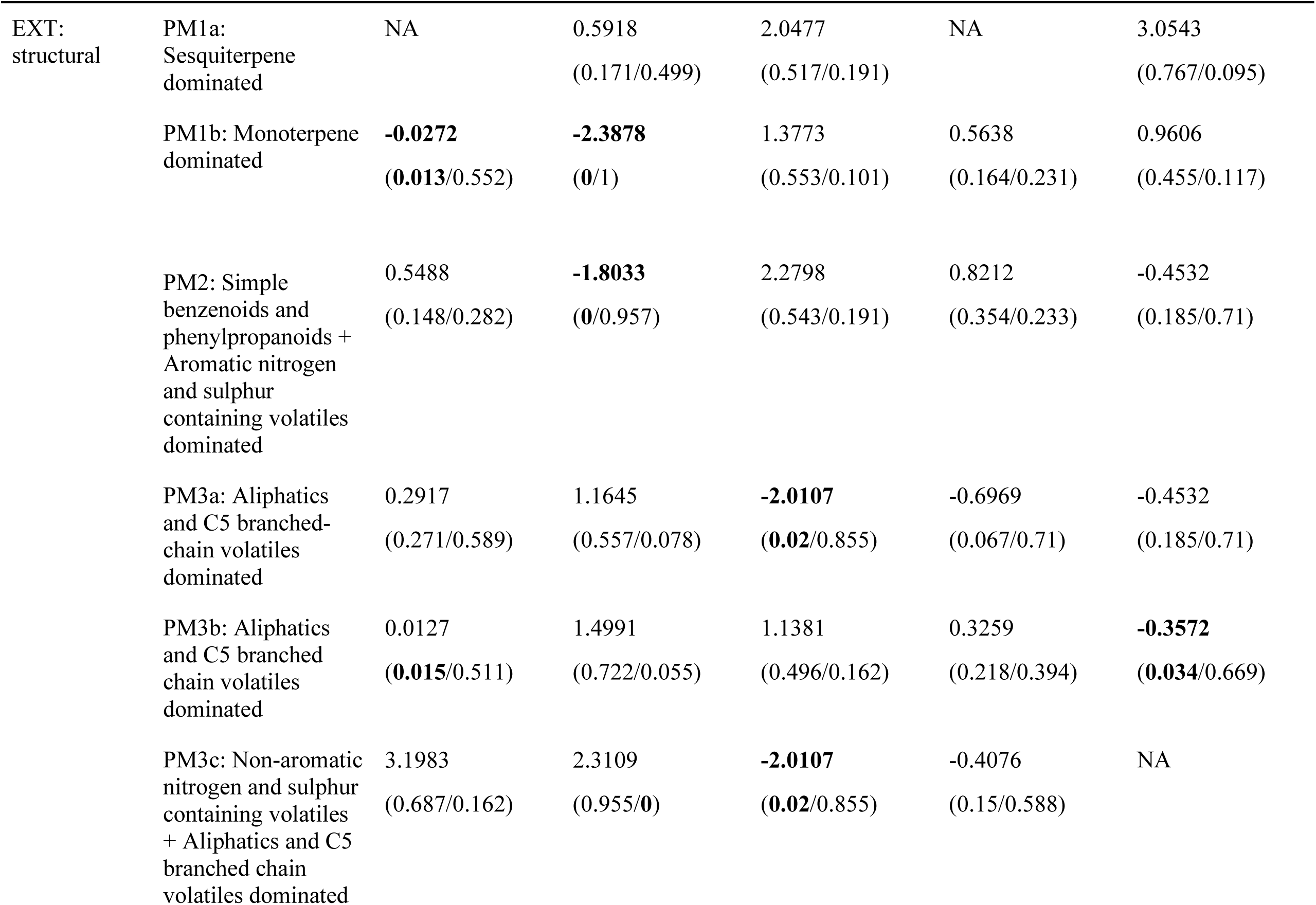
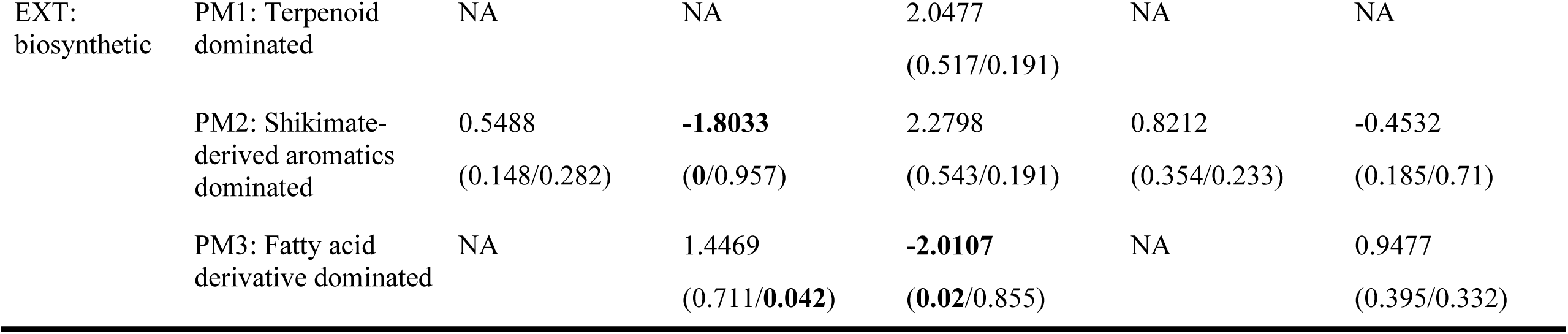
Phylogenetic signals calculated as D-statistics for EXT fingerprints across different taxonomic ranks. A value of D = 0 means phylogenetically conserved as expected under Brownian threshold motion, while D = 1 indicates random distribution of the trait across the phylogeny. Values of D < 0 indicate stronger phylogenetic clustering (traits more conserved than expected under Brownian motion), while values of D > 1 mean phylogenetic overdispersion (traits more divergent than random). p < 0.05 means the D-value is statistically significant and is represented in bold. p_r_ = Probability of estimated D resulting from no (random) phylogenetic structure; p_b_ = Probability of estimated D resulting from Brownian phylogenetic structure; PM = Plant metabolite cluster

## DISCUSSION

Plant volatiles are often considered highly labile^[32–33]^, as their emissions can be expected to vary with ecological interactions^[2]^ and the environment^[3–4]^. This plasticity in plant volatiles is suggestive of a weak phylogenetic signal in fragrance profiles, potentially limiting their taxonomic utility. However, in taxonomy, plant metabolites have often been used to differentiate populations^[14]^, species^[16]^, genera^[15,17]^, which contradicts the idea presented above that plant volatiles are labile and therefore taxonomically uninformative. One of the primary reasons for the presence of contradictions in the utility of chemical signatures as taxonomic characters is that chemical traits or features are not easily comparable, as homology between chemical compounds is challenging. Thus, chemotaxonomic studies using structural features of metabolites^[15,25,26]^, are very limited. Our study fills this knowledge gap, and we propose a machine-learning approach to assess the presence of taxonomic signatures in plant volatiles. We analysed the structural features of volatiles from curated plant volatile databases and used trait mapping to test whether these features and their associations with biosynthetic pathways are conserved within plant lineages.

Detecting taxonomic signals across diverse plant lineages requires approaches that can capture subtle high-dimensional patterns; hence, we employed data-driven learning methods on an extensive plant volatile dataset. Our results show that the ML classification models accurately predicted most plant species to their respective taxonomic ranks, indicating that plant species possess distinctive volatile profiles (Supplementary Figs. S8-S11). The inclusion of fingerprints representing structural and biosynthetic information indeed enhanced the model’s performance (Table 1), with the EXT fingerprint achieving slightly better performance, a richer, higher-bit-size type^[34]^ fingerprint, compared to fingerprints that summarised structural (SubCount) or biosynthetic (PubChem) classes. Thus, information-dense molecular fingerprints improved taxonomic assignment, enabling better discrimination among plant species. In contrast to previous ML approaches focused on a single fingerprint type^[25]^ or structural features of a single major compound class^[26]^, our method compares multiple fingerprint representations, enabling the detection of taxonomic signals across diverse plant lineages.

The volatile profile distinctiveness that distinguished plant species detected by ML likely reflects two underlying evolutionary processes acting on the volatile metabolites and their structural features: a) ecological interactions and b) phylogenetic constraints. Ecologically, pollinators and herbivores respond to complex blends rather than individual volatiles, implying selective pressures on volatile combinations^[35–36]^. Consistent with this, our models revealed that taxonomic groups are identified by distinct combinations of shared volatiles rather than by unique diagnostic compounds (Supplementary Figs. S12-S15).

Since fingerprints were found to be useful for taxonomic identification, we next explored whether the multivariate volatile profiles and fingerprint-derived volatile clusters exhibit phylogenetic signals within the ITS-based phylogeny of ML-classified plant species. First, using presence-absence data, we detected a weak but significant relationship between genetic relatedness and volatile similarity across plant species (Mantel’s test; Supplementary Note S7, Supplementary Fig. S18). Similar weak signals have been observed in studies examining broad chemical classes (such as terpenes)^[19–20]^. Our results corroborate these findings by showing that closely related species remain chemically similar, even when considering diverse volatile blends spanning multiple chemical classes.

Second, we mapped the structural and biosynthetic volatile clusters derived from the ML-fingerprinted dataset onto the phylogeny to explore phylogenetic signals in volatile clusters across taxonomic clades. Despite an uneven taxonomic sampling within the plant volatile database that we used, all three biosynthetic clusters were observed to be ubiquitous across plant orders and plant families (Fig. 4). In contrast, we identified clade-specific combinations of the six structural clusters, indicating the presence of phylogenetic structuring in the distribution of volatiles across seed plants (Fig. 3). This could mean that while broad biosynthetic classes may be widely shared across plant lineages, the distribution of structural classes might be a result of lineage-specific branching and downstream modifications of the shared metabolite precursors.

We noted that a strong phylogenetic clustering (D < 0) was recovered in three distinct structural volatile clusters in plant orders, including Myrtales (eucalyptus, guava, clove), Lamiales (mint, jasmine, basil), and Rosales (roses, apples, figs) and the higher clade Malvids (Figs. 3, 4, Table 2). At lower taxonomic ranks (such as within families), the strength of phylogenetic signals in volatile clusters varied, consistent with a taxonomic hierarchy in the distribution of volatile clusters across clades, such as the monoterpene-dominated cluster in Lamiceae and Moraceae and the aliphatic-dominated cluster in Rosaceae. The patterns likely reflect evolutionary conservation of biosynthetic pathways or gene family expansions characteristic of these lineages. For instance, the sesquiterpene-rich profiles of Lamiaceae (a major family in Lamiales)^[37]^, previously shown to exhibit strong phylogenetic constraints^[38]^, were recovered to be tightly clustered in our study, although the D-statistic could not be estimated on a single state (presence of the sesquiterpene-dominated cluster in all taxa) for Lamiaceae. Similarly, we identified clusters dominated by benzenoid, phenylpropanoid, and aromatic nitrogen- and sulphur-containing volatiles that were phylogenetically clustered within Rosales. Given the prevalence of fragrant flowers and fruits in Rosales, this strong phylogenetic clustering is consistent with the possibility that specific shikimate-derived benzenoids and aromatic nitrogen- and sulfur-volatiles, which are major components of fragrances^[39]^, may have been selected for pollinator interactions and seed dispersals in this lineage. Notably, these clades that exhibited strong phylogenetic signals also showed high classification accuracy in the ML models (Supplementary Fig. S8), suggesting that conserved volatile chemistry likely contributes to their chemotaxonomic distinctiveness.

In addition to phylogenetic clustering, phylogenetic overdispersion and randomness (D greater than or equal to 1) were also detected in our study. Phylogenetic overdispersion and randomness can arise from chemical convergence among distantly related species and from divergence among close relatives. In this study, because sampling bias can affect the presence or absence of phylogenetic signal, we refrain from drawing conclusions for clades, clusters, or taxonomic groups where taxonomic sampling was limited (< 20 tips). However, the inclusion of these taxa in the current study was deemed important, as it provides a birds-eye view of comparative volatile analysis within an evolutionary framework and adds polarity to volatile traits within the phylogenetic context. It is also important to note that the chemical similarity among species does not always imply homologous biosynthetic origins, as the same compound can be derived from multiple origins^[40–41]^. Therefore, information on enzymes involved in biosynthetic pathways might also be required to make stronger evolutionary interpretations. We hope that our study highlights opportunities for future studies to carry out targeted sampling of volatiles across different lineages to improve our understanding of the evolution of volatiles and their structural features across the angiosperm phylogeny.

In conclusion, we confirm that chemotaxonomy, which has long been suggested to be an important part of plant identification and taxonomy, is validated by the results reported in our study. While data generation and analyses are tedious, one of the most challenging aspects of using volatile profiles for phylogenetic analyses or even clustering has been the generation of characters from chemical profiles. The second challenging aspect of using metabolites as taxonomic signatures is identifying features suitable for downstream analyses. We show that fingerprint data can be utilised to infer features using machine learning models, and these validated features can then be tested using phylogenetic methods to obtain taxonomic information. We propose that advanced modelling approaches, such as graph neural networks that incorporate detailed structural features, may further improve predictive performance. While it is a common observation that botanists often crush leaves or smell flowers to obtain taxonomic identification in the field, our results empirically validate that plant volatiles carry this taxonomic information.

## METHODS

### Plant species-volatiles dataset

To identify evolutionary patterns in plant volatiles, we conducted an extensive literature review using two primary databases on plant volatiles. The databases include phytochemicals analysed and detected using techniques such as Gas chromatography (GC) coupled with Flame Ionisation Detector (FID) or Mass spectrometry (MS). The AROMA DB (http://bioinfo.cimap.res.in/aromadb/)^[42]^, developed by the Council of Scientific & Industrial Research-Central Institute of Medicinal and Aromatic Plants (CSIR-CIMAP), is a comprehensive electronic repository that catalogues information on 1322 volatiles sourced from nearly 190 medicinal and aromatic plant species of both Indian and foreign origins. The Pherobase database (https://www.pherobase.com/)^[43]^ specializes in pheromones and semiochemicals and includes data on floral volatiles from approximately 101 plant families. We manually compiled the training dataset from the literature retrieved through the two databases above, which included species-volatile information for 70 plant families, comprising 66 angiosperms and 4 gymnosperms, resulting in a total of 691 plant samples. Taxonomic annotation was performed at the order, family, and genus ranks.

Unidentified volatiles or generic labels (e.g., sesquiterpene 1) were omitted because their m/z values were not consistently mentioned across different studies. Volatiles in trace quantities were included to obtain a near-complete volatile profile for each species. We encoded the dataset in a binary format (1 = detected in any sample of the species, 0 = absent) because quantitative information has been reported inconsistently across different studies (relative percentages, absolute concentrations, or maximum emissions of volatiles).

### Volatiles-molecular fingerprint dataset

To retrieve the structural details of the volatiles synthesised by various plant species, we first determined the Simplified Molecular Input Line Entry System (SMILES) notations for each of the volatiles. This format provides a one-dimensional representation in which a compound is represented as a simple sequence of characters with predefined atom-ordering rules^[44]^. SMILES notations for volatiles were sourced from the PubChem compound database (https://pubchem.ncbi.nlm.nih.gov/)^[45]^. After consolidating the SMILES notations, the input data were uploaded to the PaDEL descriptor software^[34]^ to generate different types of substructure-based molecular fingerprints. The fingerprint types included both binary (PubChem - 881 bits, MACCS - 166 bits, Sub - 304 bits, and EXT - 1024 bits) and count (SubCount - 304 bits) types.

### Clustering of plant volatiles using fingerprint data

To determine the optimal clusters from each fingerprint type, clustering techniques based on distance and density were applied to the volatiles-molecular fingerprint dataset for five different fingerprint types. Distance-based methods included k-means and agglomerative clustering, whereas the density-based approach was DBSCAN. We also used spectral clustering, a hybrid clustering approach that leverages graph theory and eigenvalues.

For performance comparison, we calculated the Silhouette score, Davies-Bouldin score, and Calinski-Harabasz score, where the Silhouette score measures how similar a sample is to its own cluster compared to other clusters, the Davies-Bouldin score measures the average similarity between each cluster and its most similar one, and the Calinski-Harabasz score indicates the ratio of between-cluster dispersion to within-cluster dispersion. The Calinski-Harabasz score showed a stronger correlation with the cluster plots (Supplementary Table S2); hence, we will discuss this score in further analyses. In the Calinski-Harabasz score, a higher score indicates better clustering. To visualise high-dimensional patterns and inspect cluster coherence, we used t-distributed stochastic neighbour embedding (t-SNE), which performs a non-linear projection of the data, preserving local structure and providing a better representation than other ordination methods^[46]^.

### Cluster validation

We expected that structurally similar clusters might be linked to major biosynthetic origins, especially at broad class levels^[25]^. To determine whether the resultant volatile clusters reflected structural and biosynthetic features, all volatiles were categorised into two groups, namely, structural classes^[47]^ and broad biosynthetic classes^[39,48]^, respectively. The structural classes were based on their core backbones and functional groups. They included eight classes, simple benzenoids and phenyl propanoids (aromatic benzene rings with often C6-C3 structures), aromatic nitrogen and sulphur containing volatiles (aromatic rings with nitrogen, sulphur or both), non-aromatic nitrogen and sulphur containing volatiles (no aromatic rings, but has nitrogen, sulphur or both), monoterpenoids (cyclic or acyclic C10 volatiles derived from isoprene units), sesquiterpenoids (C15 volatiles with complex rings, also derived from isoprene units), diterpenoids (C20 volatiles from isoprene units), irregular terpenoids (do not follow strict C5 rule, but still derived from isoprene units) and aliphatics and C5 branched chain volatiles (straight chain or branched saturated and unsaturated volatiles; with no aromatic rings). Biosynthetic classes were based on their major pathways (Supplementary Fig. S4), which included shikimate-derived aromatics derived from the shikimate-phenyl propanoid pathway, terpenoids synthesised from the mevalonate (MVA) or methylerythritol phosphate (MEP) pathways, fatty acid derivatives including aliphatics derived from the lipoxygenase (LOX) pathway, and remaining volatiles with mixed or ambiguous origins classified as others.

The alignment of the obtained clusters to the manually annotated volatile classes was estimated using the normalised mutual information (NMI) score, which ranged from 0 to 1, and the adjusted rand index (ARI), which ranged from -1 to 1; in both cases, higher values indicated better overlap between discovered and ground-truth clusters. The NMI and ARI scores were estimated using the “aricode” and “mclust” packages in R version 4.5.1^[49]^, respectively. The optimal number of clusters was determined based on these scores and t-SNE plots.

### Fingerprint inclusion and feature selection for classification

To predict plant species at the taxonomic ranks of order, family, and genus, we trained four different classification models in two ways: (1) binary plant volatiles data alone and (2) the same data combined with each fingerprint type. We examined whether the fingerprint type that achieved the highest NMI/ARI scores in the clustering analysis discussed in the previous section also yielded the best classification performance.

To achieve this, we combined the fingerprint data with the plant volatiles as a whole, independent of any clustering methods used in previous studies^[25–26]^. For a data point, we filtered all volatiles from the fingerprint dataset, then computed the mean fingerprint for each set of volatiles, and combined the results with the original volatile data. For feature selection, we applied a three-step process that includes Jaccard similarity-based selection, correlation-based selection, and filtering out near-constant features (for a detailed description of the filtering steps, refer to Supplementary Note S1 and Supplementary Table S4). These steps collectively reduced dimensionality, improved computational efficiency, and ensured that only informative and non-redundant features were retained for subsequent modelling.

### Model training and evaluation

We employed four supervised machine learning models, including logistic regression (LR), random forest (RF), support vector classifier (SVC), and multilayer perceptron (MLP). For a detailed description of classification models, refer to Supplementary Note S2.

We adopted a two-step analysis to train the models. In the first step, we used a grid search to identify the optimal model by tuning the hyperparameters. The grid search systematically explored a predefined set of values for each hyperparameter, training and validating the model on randomly selected subsets of the data. We partitioned the dataset into 80% for training and 20% for testing, and selected the best-performing hyperparameters based on validation. In the second step, we trained and evaluated the model using 5-fold cross-validation to ensure robustness and reduce the risk of overfitting. In this procedure, the data were divided into five equal parts, with each fold (20% of the data) serving once as validation, while the remaining folds were used for training. The final test results were derived from the aggregated performance across all five folds and the average of five runs (the final modelling was performed five times), providing a more reliable estimate of the model’s generalisation ability.

The prediction performance of the test data was evaluated using metrics such as accuracy, precision, recall, and F1 score. A weighted average for all metrics was used to account for the unbalanced dataset. Confusion matrices were generated to identify plant species that were correctly predicted to true taxonomic ranks or misclassified into different ranks. Higher values in the diagonals of the matrix represent a perfect match.

### Identifying key features for the prediction of taxonomic ranks

To identify the important features, we used permutation scores and model-specific methods across different algorithms, given their differences in working principles. Here, the permutation score calculates the feature importance by checking the extent to which a model’s performance metric decreases when the values of a single feature are randomly shuffled (permuted). Hence, if shuffling a feature causes a significant drop in performance, it implies that the feature is important, and if shuffling makes little or no difference, then the feature is not important, as the model does not rely on it. In the case of LR, we identified important features based on model coefficients; in the case of RF, we used Gini; and for the MLP, we used only permutation scores due to the model’s black-box nature. **Mapping of volatile clusters and calculation of phylogenetic signals**

To explore the presence of phylogenetic clustering in the distribution of plant volatiles, we mapped volatile clusters (refer to the “Clustering of plant volatiles using fingerprint data” section above) from selected fingerprint types onto a phylogenetic tree (for tree building protocols, see Supplementary Note S6). We used the volatile clusters from the fingerprint type that showed the best classification performance across taxonomic ranks. The optimal clusters to map, representing structural and biosynthetic classes, were identified based on NMI/ARI scores. Volatile clusters were coded as 1s and 0s, where 1 represents the presence of any volatile from that particular cluster in the plant species, and 0 represents its absence. Trait mapping and visualisation were performed using Interactive Tree Of Life (iTOL; version 7.2.2)^[50]^. Because the traits were represented in binary format, we used the Fritz-Purvis D-statistics to estimate phylogenetic signals^[51]^ with significance assessed under Brownian motion or random shuffle (based on 1000 permutations), as implemented in the phylo.d function of the “caper” package in R version 4.5.1^[49]^. The D-statistic was calculated similarly to the method described by Zhang et al.^[36]^ for the entire phylogeny and for the 11 major clades identified in the phylogenetic tree (that included Magnoliids, Commelinids, Rosids, Asterids, Fabids, Malvids, Lamiids, Campanulids, Rosales, Myrtales, and Lamiales), with at least 20 tips and at least three families represented. The D-statistic was also calculated for the families, including Lamiaceae, Asteraceae, Apiaceae, Moraceae, Rosaceae, and Myrtaceae, that dominated these major clades.

## Data availability

All datasets and codes generated in this study will be made available in a public repository once the study is published.

## Funding

The computational part of this work was funded from the Science and Engineering Research Board (SERB POWER grant: SPG/2021/000793), awarded to V.G. and A.D. V.G. discloses support for the research of this work from the Ministry of Human Resource Development (Ministry of Education). A.S. discloses support for the timely funding of the fellowship from the Department of Biotechnology (DBT/2020/IISER-B/1309).

## Acknowledgement

The authors are extremely grateful to Prof. Robert A. Raguso for reviewing the first draft of this manuscript and providing valuable comments. The authors thank the Department of Biological Sciences, IISER Bhopal, and the Division of Electrical, Electronics, and Computer Sciences at IISc, for the infrastructure provided to carry out all computationally intensive analyses. The authors are extremely grateful to Abhishek, Ishanee, Ayantika, Nevil, Anjali, Antariksh, and Devika for their assistance in cross-checking the datasets. The authors also acknowledge the guidance provided for the phylogenetic work by Dr. Aleena Xavier and Dr. Ritu Yadav.

## Author contributions

V.G. and A.D. conceived the idea and designed the study. A.S. consolidated the chemical and genetic data and carried out the statistical and phylogenetic analyses. A.A. conducted the machine learning classification and clustering algorithms. A.S. and A.A. drafted the manuscript, and V.G. and A.D. edited the manuscript and gave critical comments. All authors read and approved the final manuscript.

## Competing interest

The authors have no conflict of interest to declare.

## Notes

### Competing Interest Statement

The authors have declared no competing interest.

