## Supplementary information for "Revealing taxonomic signals in plant volatiles with phytochemistry, machine learning and trait mapping"

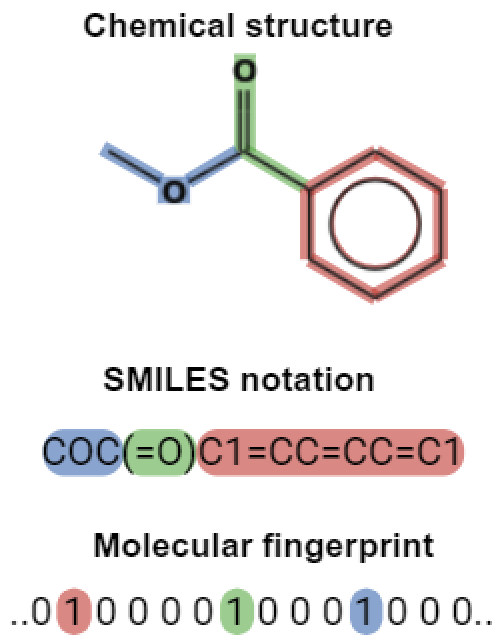


**Supplementary Fig. S1. An illustration of generation of a molecular fingerprint.** Once a metabolite is identified and SMILES notation is obtained from PubChem database, the PaDEL descriptor detects the predefined substructural features in the metabolite (denoted in different colours) and encodes their presence or absence as fingerprint bits.

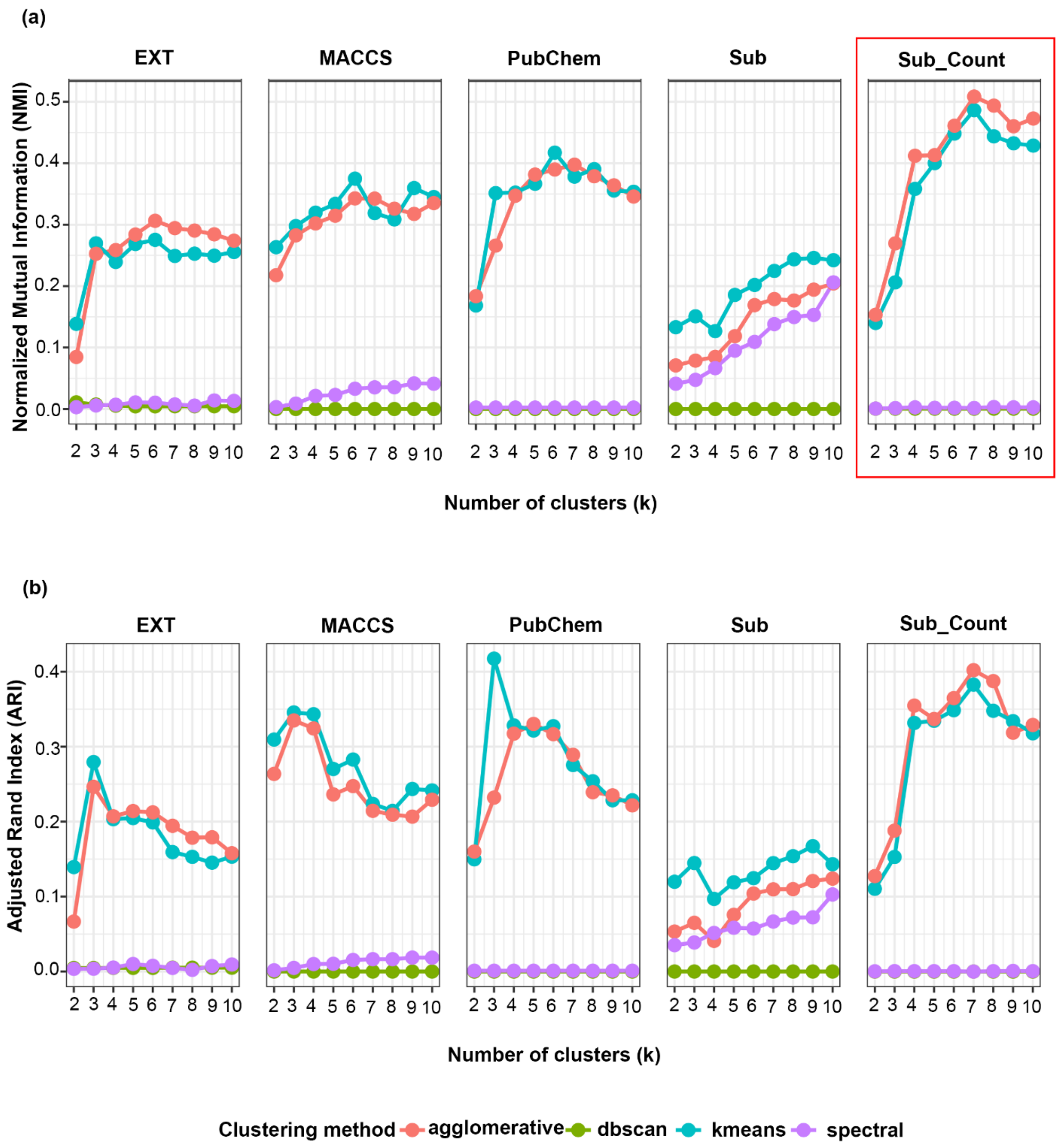


**Supplementary Fig. S2. Cluster validation across fingerprint types (structural classes).** Comparison of **(a)** NMI and **(b)** ARI scores across five fingerprint types assessing how well the predicted clusters represented ground truth labels (structural classes). Higher values mean better representation. The fingerprint type with the highest NMI score is marked with a red box. Overall, the distance-based methods (k-means and agglomerative) outperformed density-based (DBSCAN) and spectral clustering.

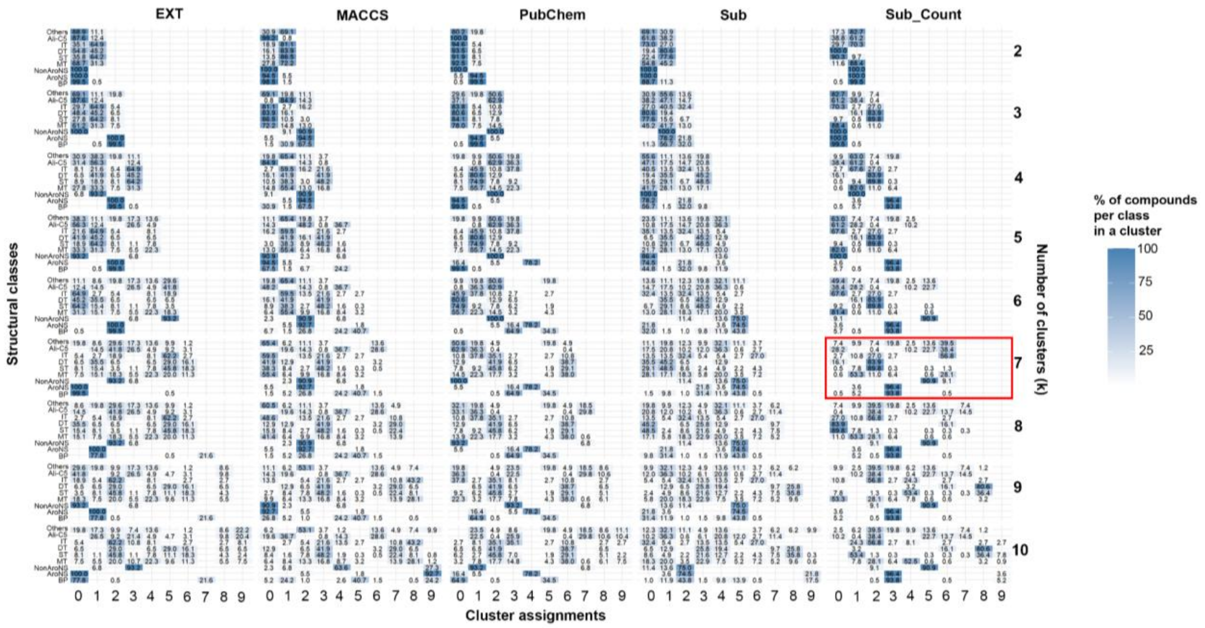


**Supplementary Fig. S3. Contingency heatmap showing volatile metabolite distribution across structural classes for clusters generated using different fingerprint types after agglomerative clustering.** The volatile metabolite distribution after agglomerative clustering of SubCount fingerprint at k=7 is represented with a red box.

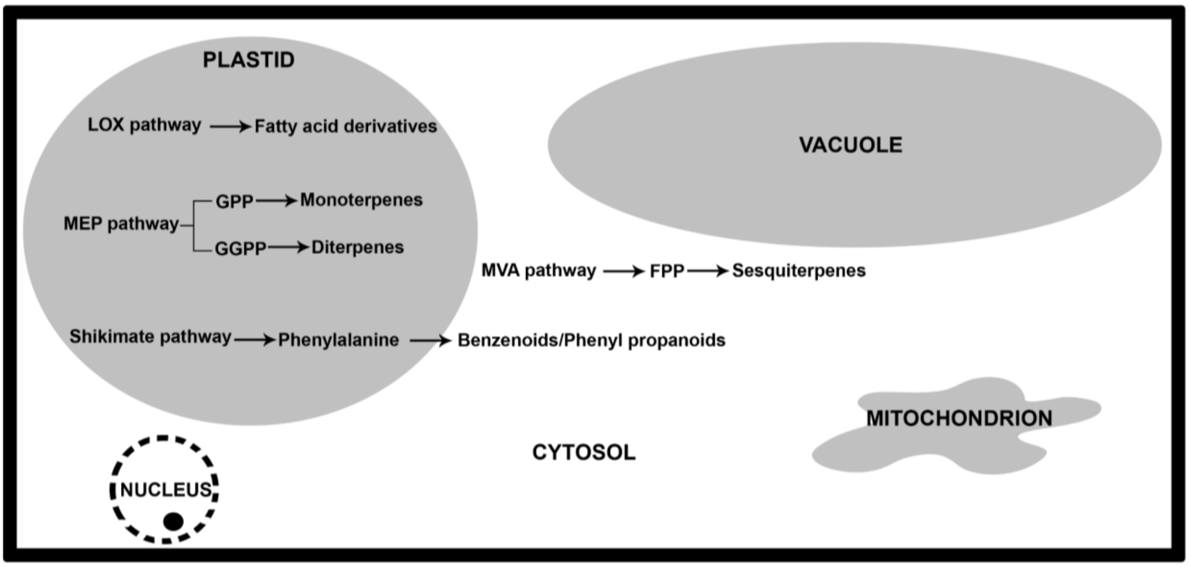


**Supplementary Fig. S4. Biosynthetic pathways for volatile metabolites in plants.** The illustration represents the synthesis of major volatile metabolite classes inside a plant cell. Figure modified from Pichersky *et al*[1] and Lv *et al*[2]. Abbreviations: GPP - Geranyl pyrophosphate; GGPP - Geranylgeranyl pyrophosphate; LOX - Lipoxygenase; MVA -Mevalonic acid; MEP - 2-methylerythritol 4-phosphate

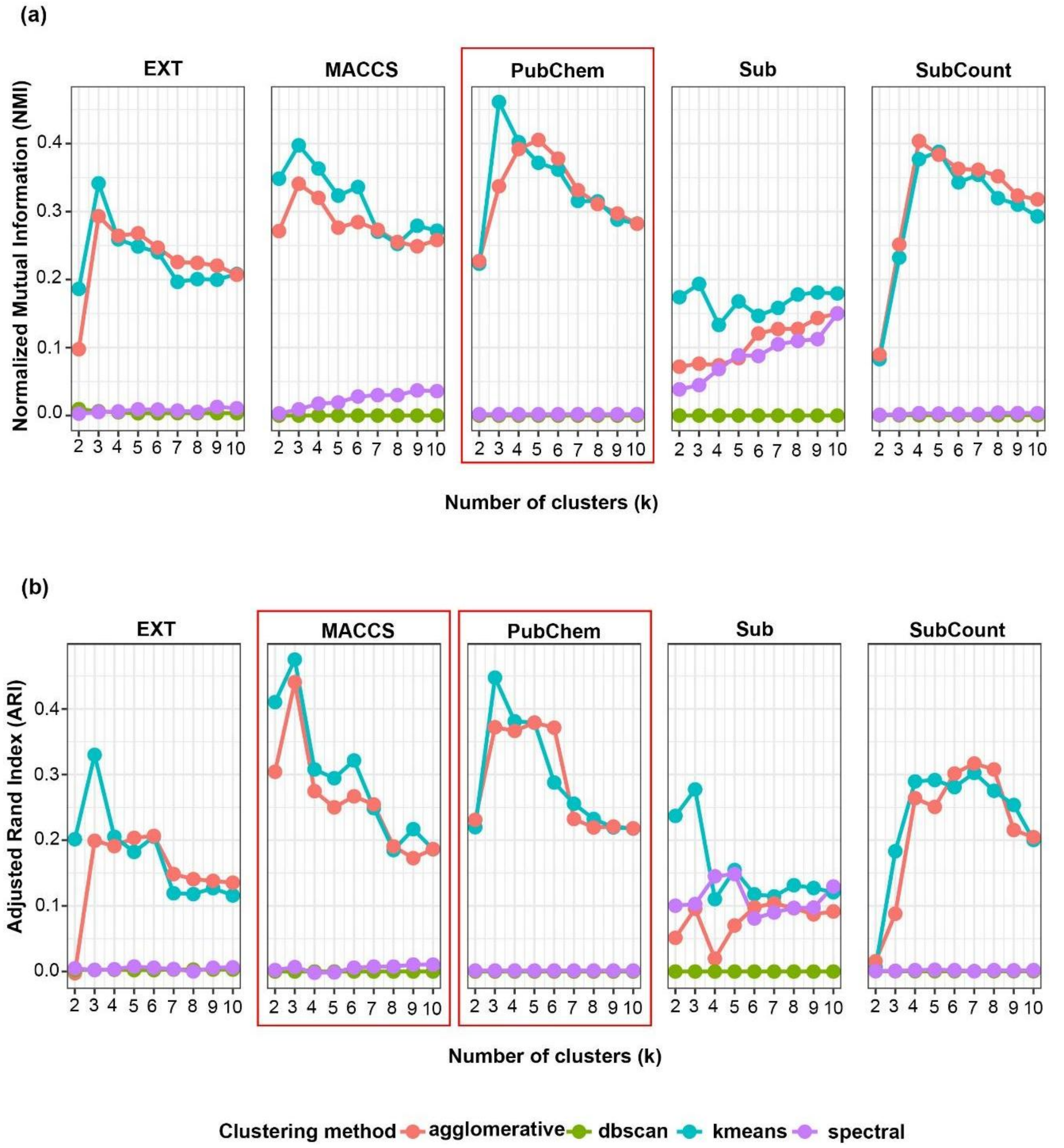


**Supplementary Fig. S5. Cluster validation across fingerprint types (biosynthetic classes)** Comparison of **(a)** NMI and **(b)** ARI scores across five fingerprint types assessing how well the predicted clusters represented ground truth labels (biosynthetic classes). Higher values mean better representation. The fingerprint types with the highest NMI score are marked with a red box. Overall, the distance-based methods (k-means and agglomerative) outperformed density-based (DBSCAN) and spectral clustering.

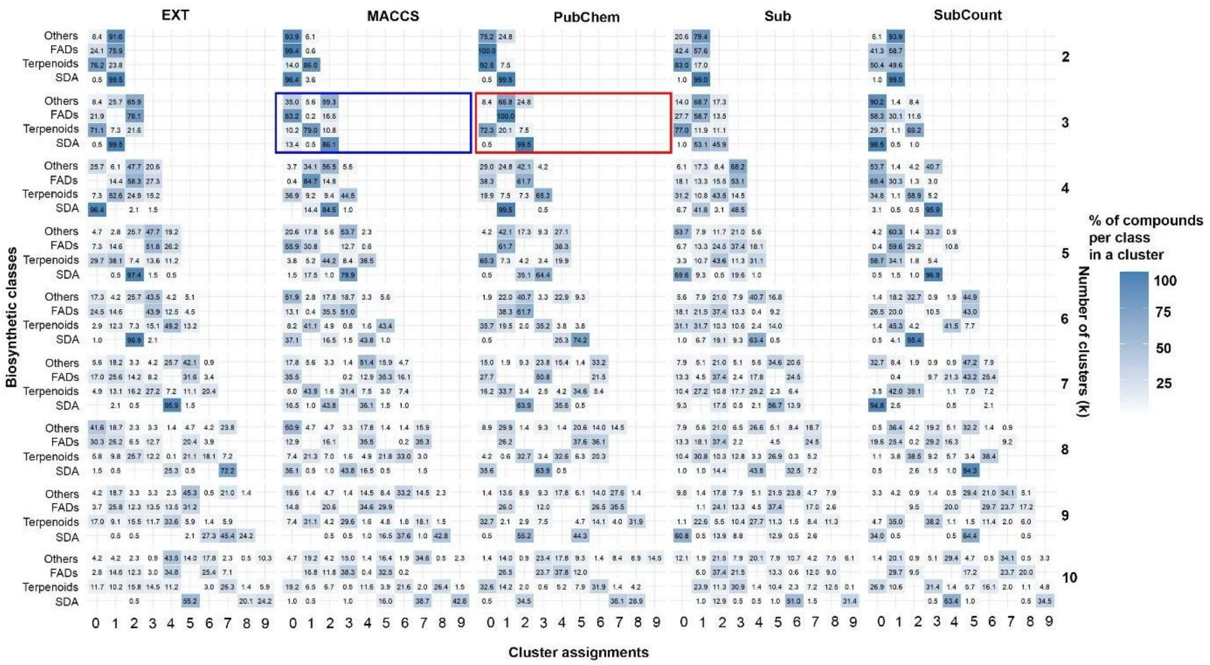


**Supplementary Fig. S6. Contingency heatmap showing volatile metabolite distribution across biosynthetic classes for clusters generated using different fingerprint types after k-means clustering.** The volatile metabolite distribution after k-means clustering of the PubChem fingerprint at k=3 is represented with a red box and that of MACCS fingerprint at k=3 is represented with a blue box.

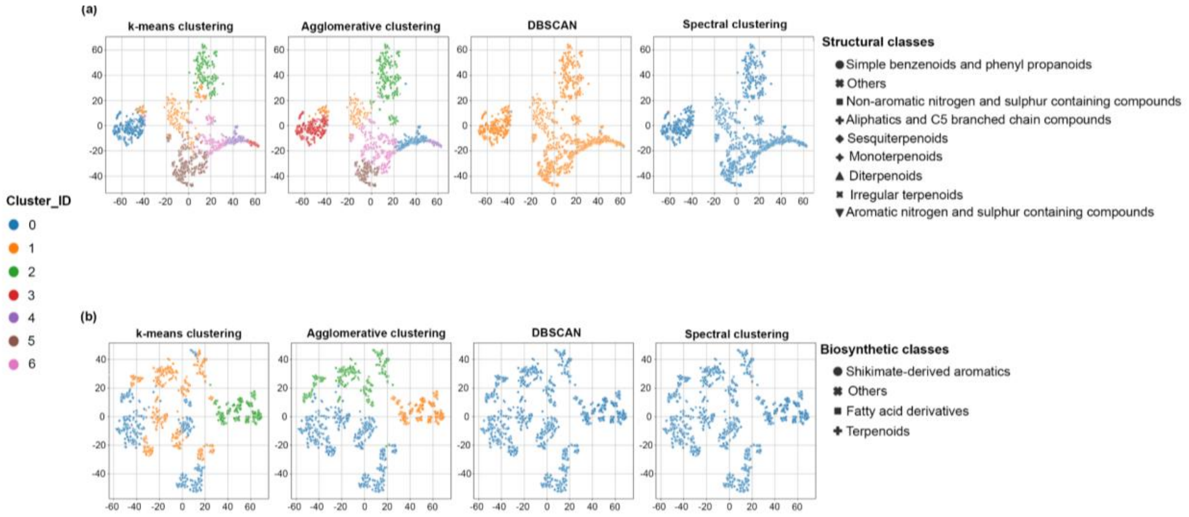


**Supplementary Fig. S7. t-SNE visualisation of clustering patterns. (a)** SubCount fingerprint type at k=7 shape-coded based on structural classes and **(b)** PubChem fingerprint type at k=3 shape-coded based on biosynthetic classes across four different clustering methods. Colours represent the cluster assignments. Overall, the distance-based methods (k-means and agglomerative) outperformed density-based (DBSCAN) and spectral clustering.

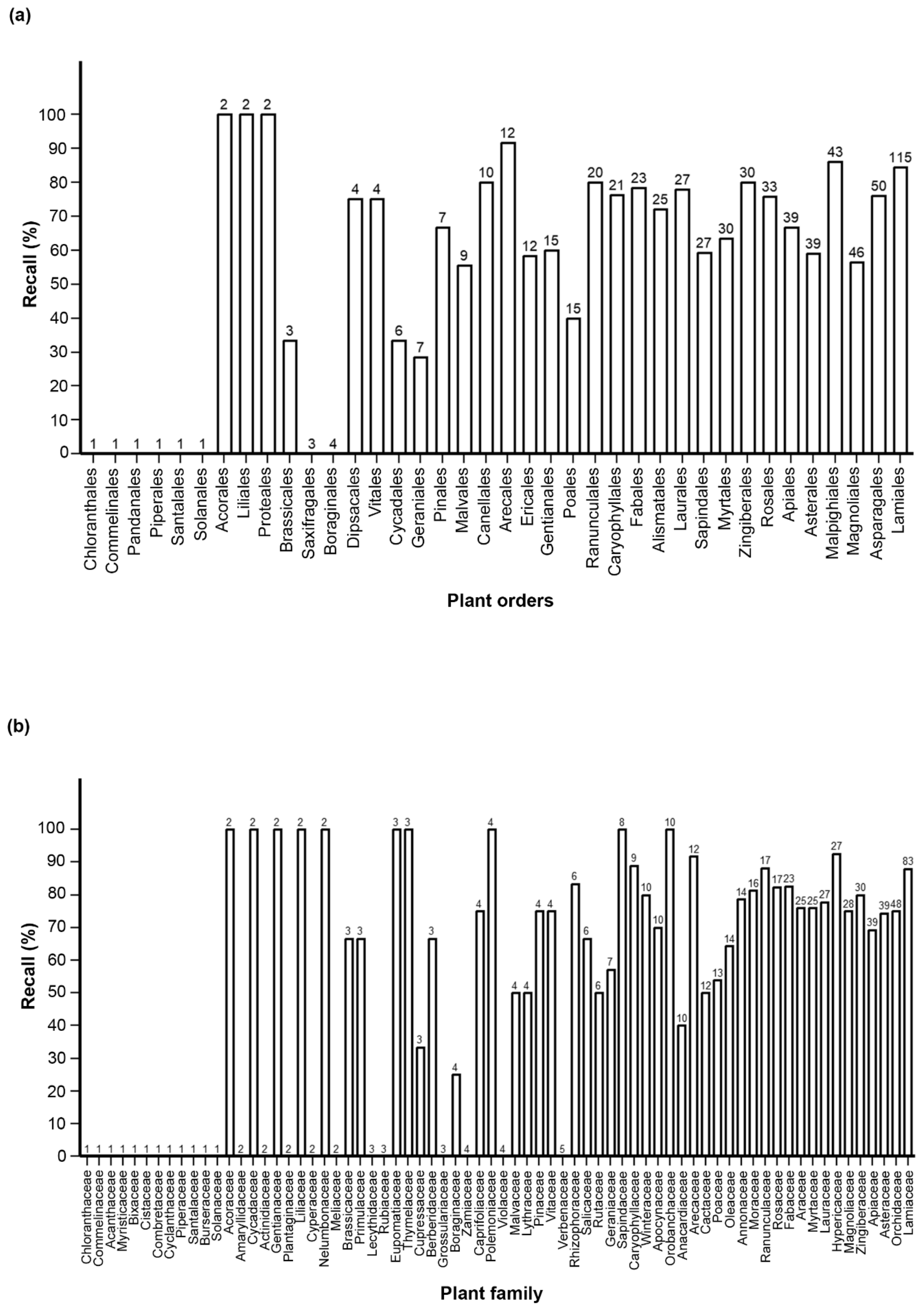


**Supplementary Fig. S8. Performance of the best classification models using EXT fingerprints across plant species.** Bar plots representing the percent of plant species accurately predicted to the right taxonomic rank in the best classification models (MLP for order and family with EXT fingerprints as features), as denoted by recall across **(a)** plant orders and **(b)** plant families. The numbers above the bars indicate the total number of data points sampled per taxonomic rank.

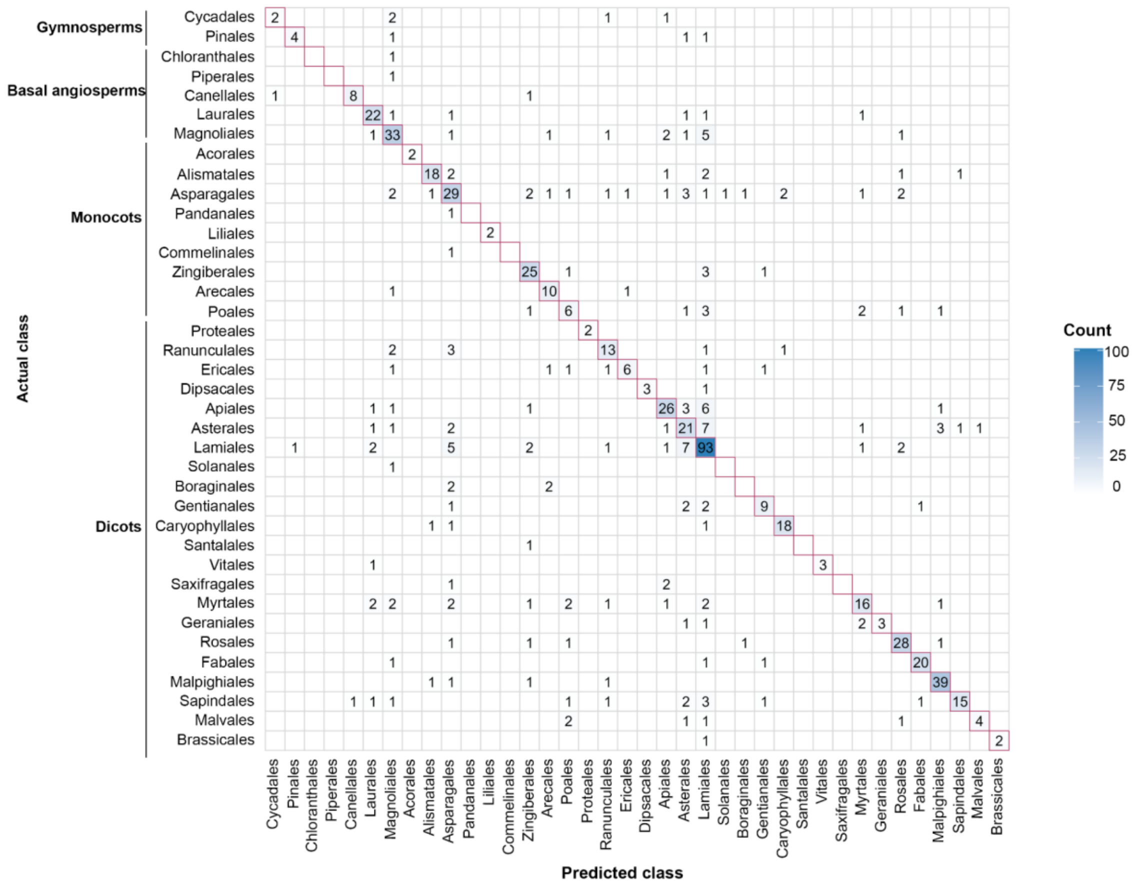

**Supplementary Fig. S9. Order-level classification performance (MLP model) with EXT fingerprints.** Confusion matrix for order-level prediction using the MLP model after cross-validation. The y-axis represents the true orders arranged according to their taxonomic relationships, and the x-axis represents the predicted orders.

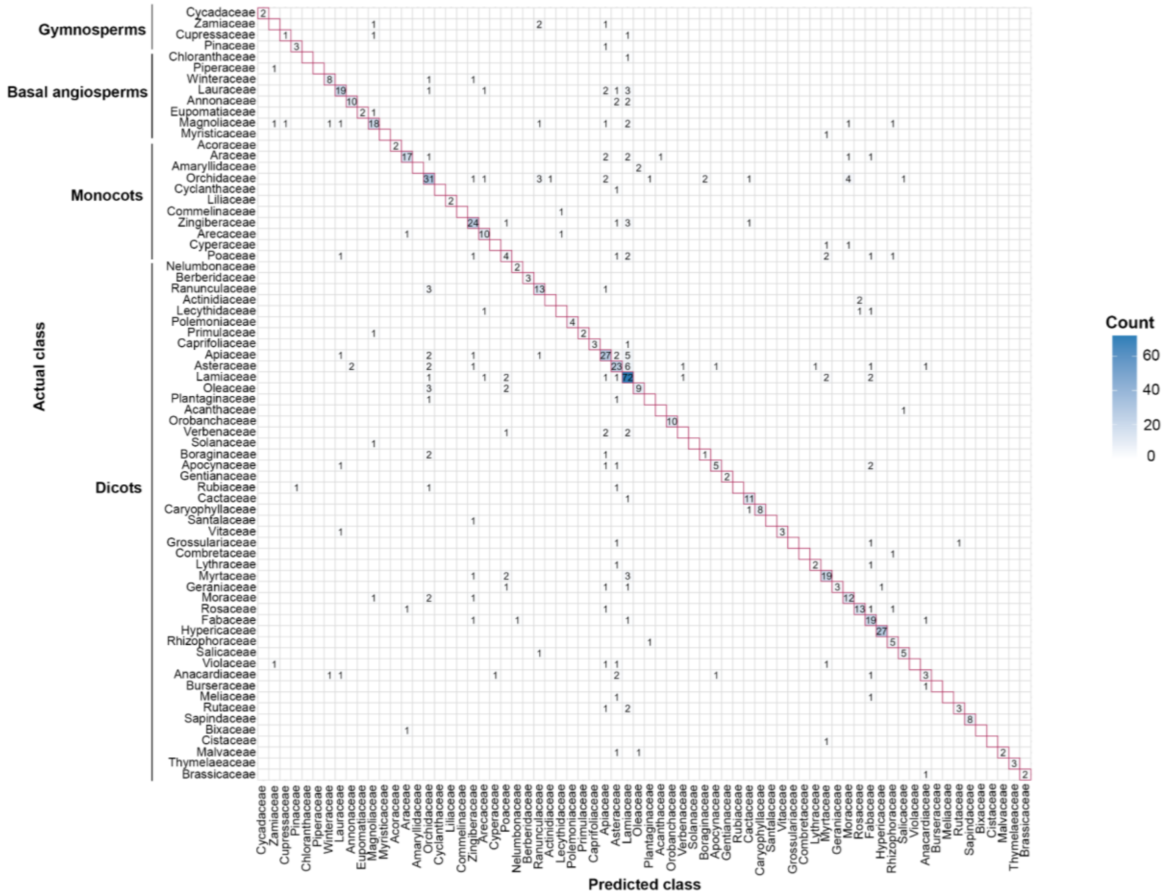


**Supplementary Fig. S10. Family-level classification performance (MLP model) with EXT fingerprints.** Confusion matrix for family-level prediction using the MLP model after cross-validation. The y-axis represents the true families arranged according to their taxonomic relationships, and the x-axis represents the predicted families.

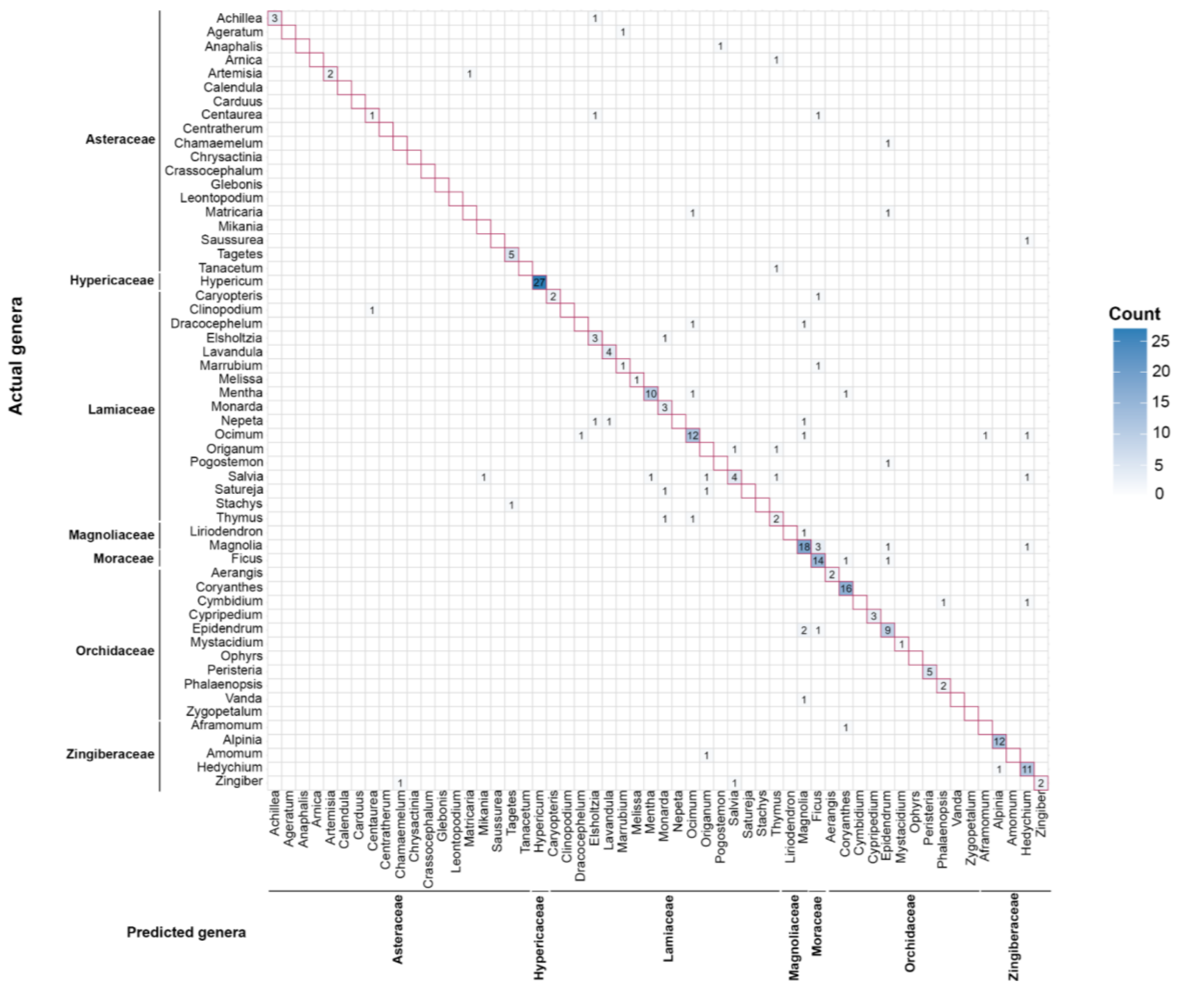


**Supplementary Fig. S11. Genus-level classification performance (LR) model with Sub fingerprints.** Confusion matrix for genus-level prediction using the LR model after cross-validation. Here, only selected families were represented that either had diverse genera or were dominated by a single genus in the training dataset. The y-axis represents the true genera arranged according to their families, and the x-axis represents the predicted genera.

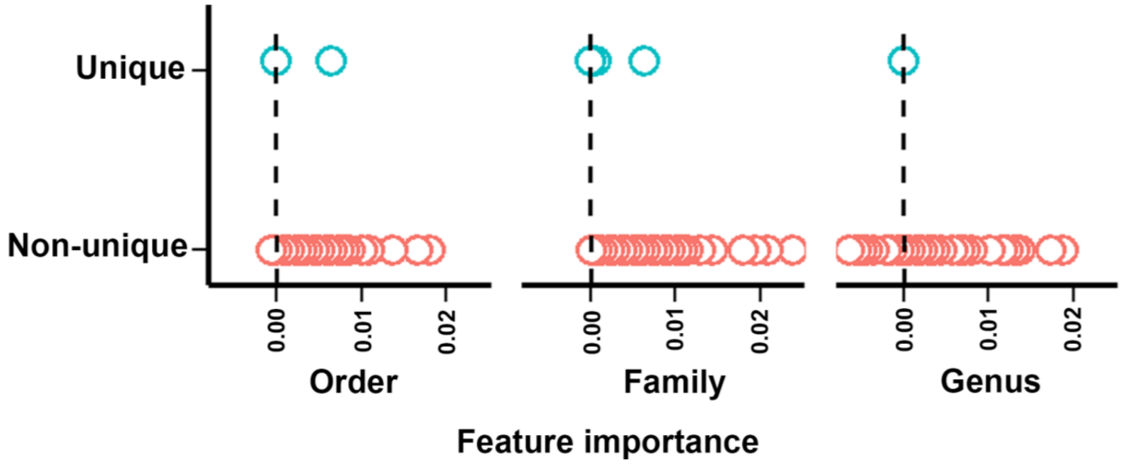


**Supplementary Fig. S12. Feature importance and taxonomic uniqueness of volatile metabolites.** Scatter plot showing permutation importance of volatile metabolites (x-axis) against their uniqueness to a taxonomic rank. Each point represents a single metabolite, with colour indicating whether it is unique (blue) or non-unique (red).

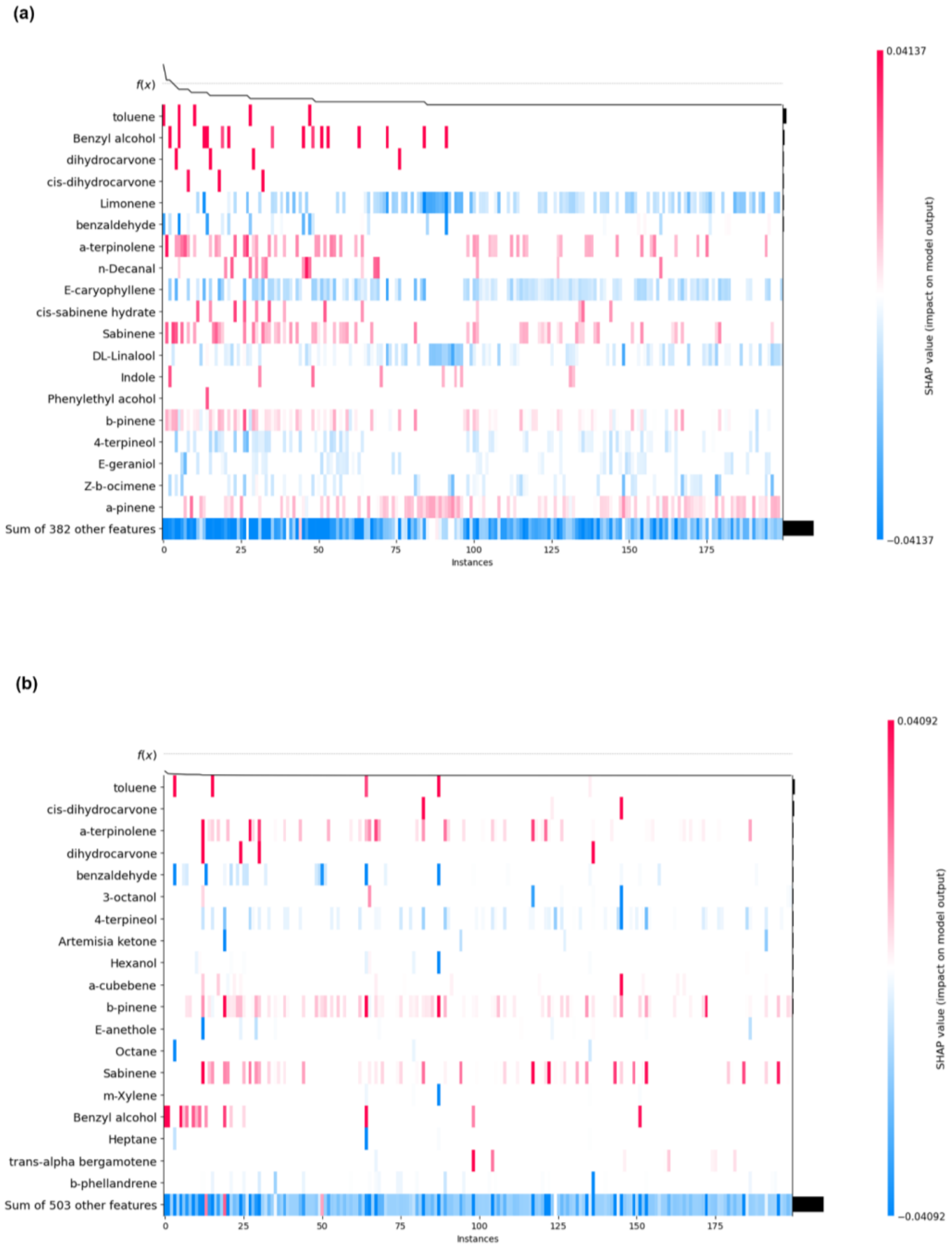


**Supplementary Fig. S13. Features determining the predictions at the order level of classification.** Shap image comparison for **(a)** volatile metabolites (RF model) versus **(b)** in combination with EXT fingerprints (MLP model). Both models strongly agree that toluene is the most critical positive predictor for classifying orders. However, the addition of EXT fingerprints dramatically simplifies the model's decision-making process. The model without fingerprints displays a "busier" and more complex logic. Lacking the specialized fingerprint data, it is forced to find a pattern by considering a much broader combination of many different compounds (like limonene, α-terpinolene, and n-decanal). This results in a "spread-out" logic where many features have a small, mixed influence. In contrast, the model with EXT fingerprints has learned a "cleaner," more focused set of rules. It relies on a few strong, clear signals (like toluene for "yes" and b-phellandrene for "no") while ignoring the noise from most other compounds. This strongly suggests the fingerprint features provide a powerful, condensed, and more direct signal, allowing the model to find a much simpler path to the correct answer.

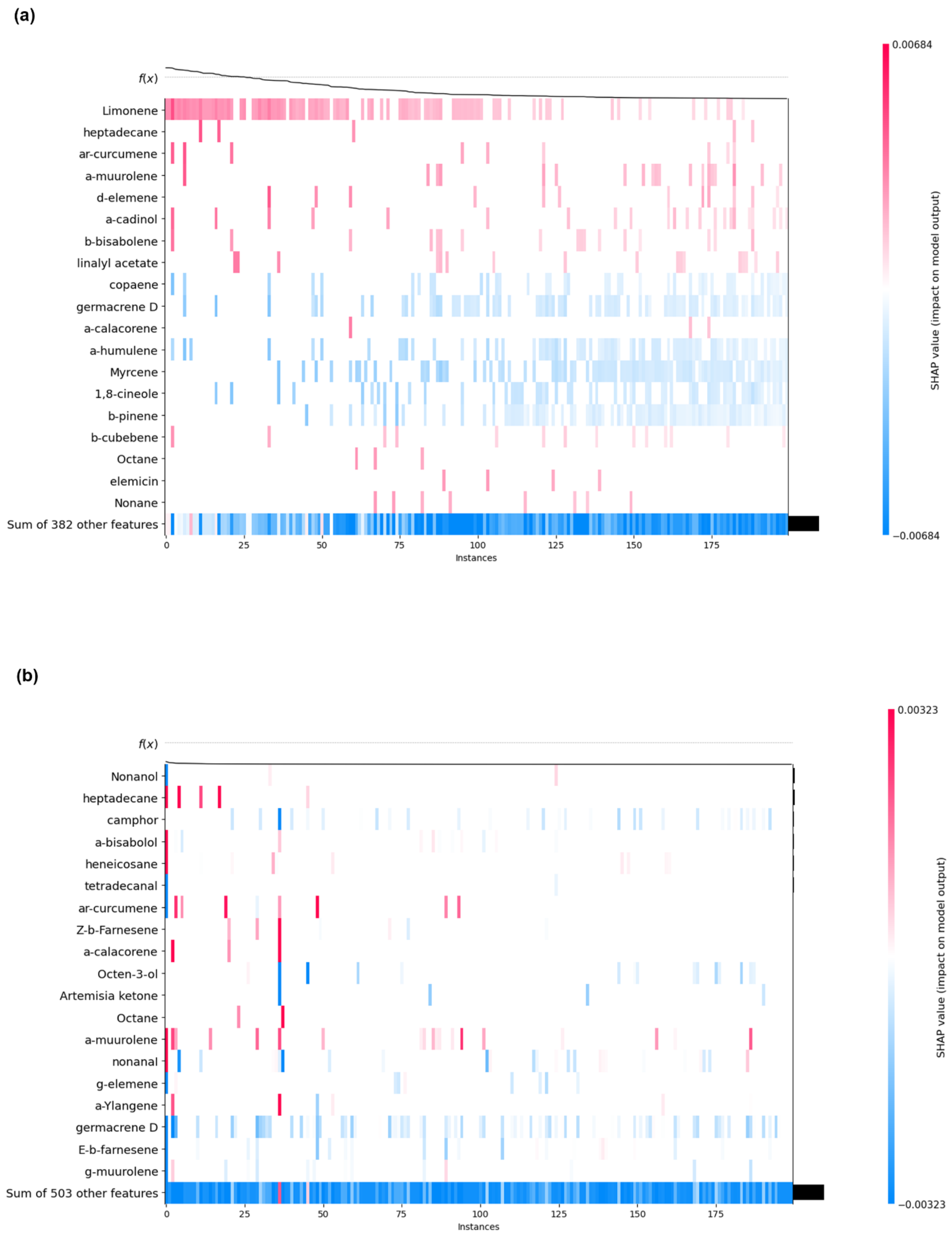


**Supplementary Fig. S14. Features determining the predictions at the family level of classification.** Shap image comparison for **(a)** volatile metabolites (LR model) versus **(b)** in combination with EXT fingerprints (MLP model). The image shows that the two models (without/with fingerprints) use two completely different strategies for identifying the families. The model without the fingerprints uses a "pro/con" logic. It relies heavily on limonene as a strong positive ("yes") signal. At the same time, it has learned a clear list of negative ("no") signals, such as germacrene D, α-humulene, and myrcene, that consistently push the prediction away from this family. The model with EXT fingerprint uses an "exception-based" logic. Its decision is dominated by the "sum of other features," which acts as a powerful default "no" vote (the solid blue bar at the bottom). The model only classifies a sample as "yes" if it finds a few specific positive exceptions to overcome this default, with heptadecane being its most important positive signal.

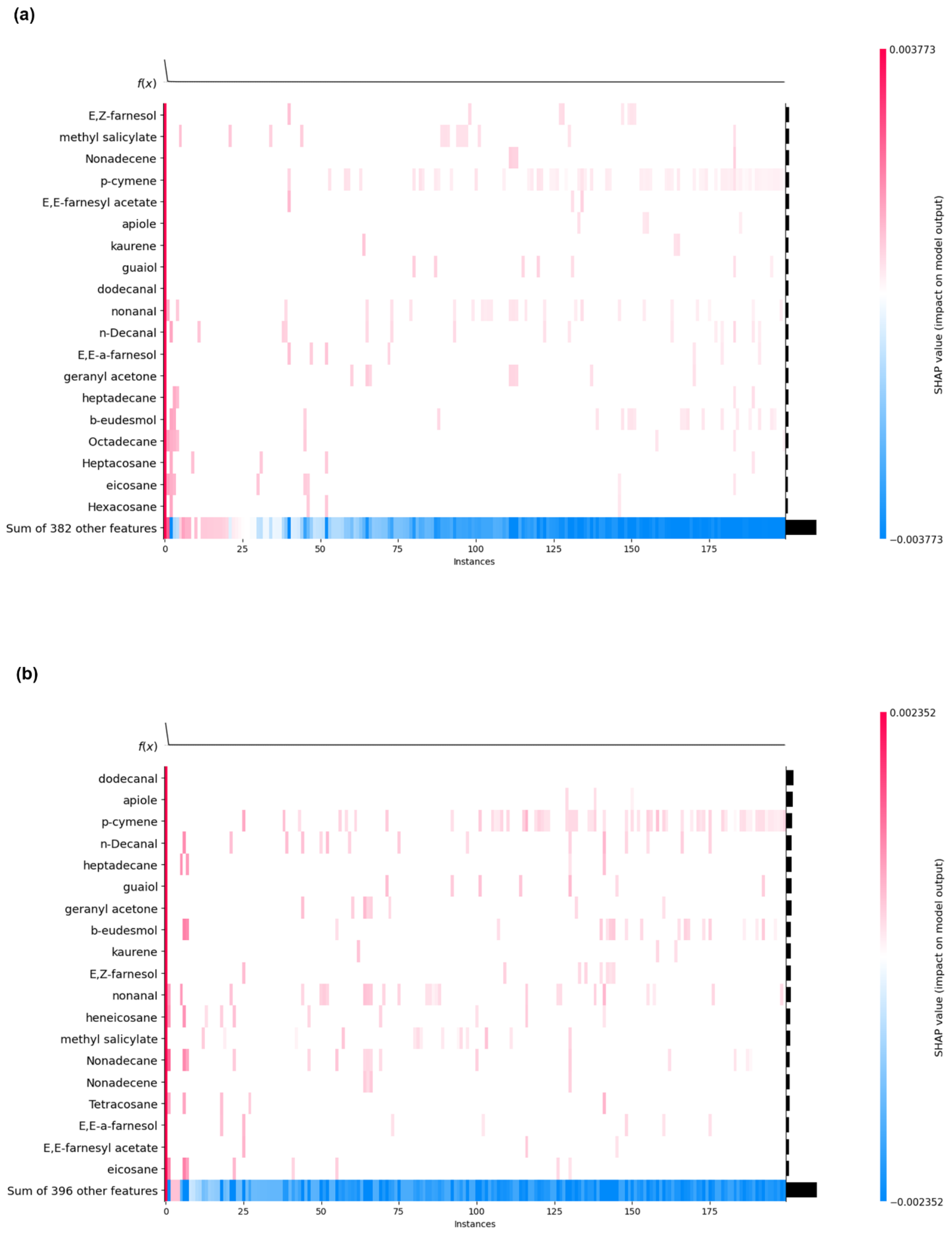


**Supplementary Fig. S15. Features determining the predictions at the genus-level of classification.** Shap image comparison for **(a)** volatile metabolites (LR model) versus **(b)** in combination with Sub fingerprints (LR model). Both models adopt a very sparse and focused logic; that is, they rely on a small handful of compounds as positive ("yes") signals for this genus, while the combined "sum of other features" acts as a strong, collective negative ("no") signal (the solid blue bar at the bottom). The model with Sub fingerprints identifies dodecanal, apiole, and p-cymene as its primary positive indicators. The model without fingerprints ignores those and instead learns that E, Z-farnesol, methyl salicylate, and nonadecene are the most important positive signals. The only notable overlap is p-cymene, which both models agree has a positive impact, though it's much more important for the model that included fingerprints.

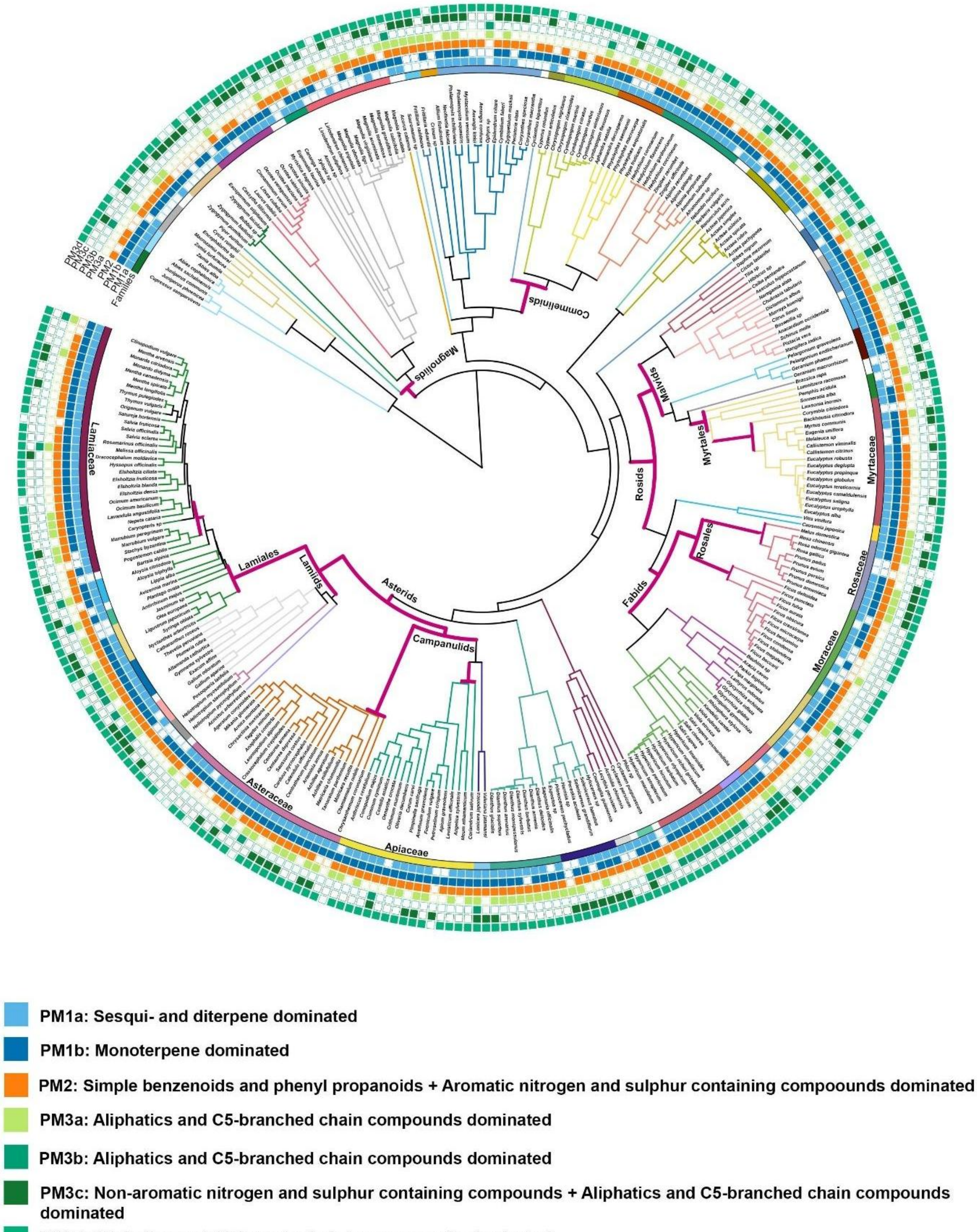


**Supplementary Fig. S16. Mapping of SubCount fingerprint-derived clusters onto the phylogeny.** Coloured branches represent the different orders and coloured stripes represents different families. Clusters are represented as coloured squares, where presence is indicated by filled squares and absence as unfilled squares. The clades analysed for D-statistics are marked as bold magenta lines.

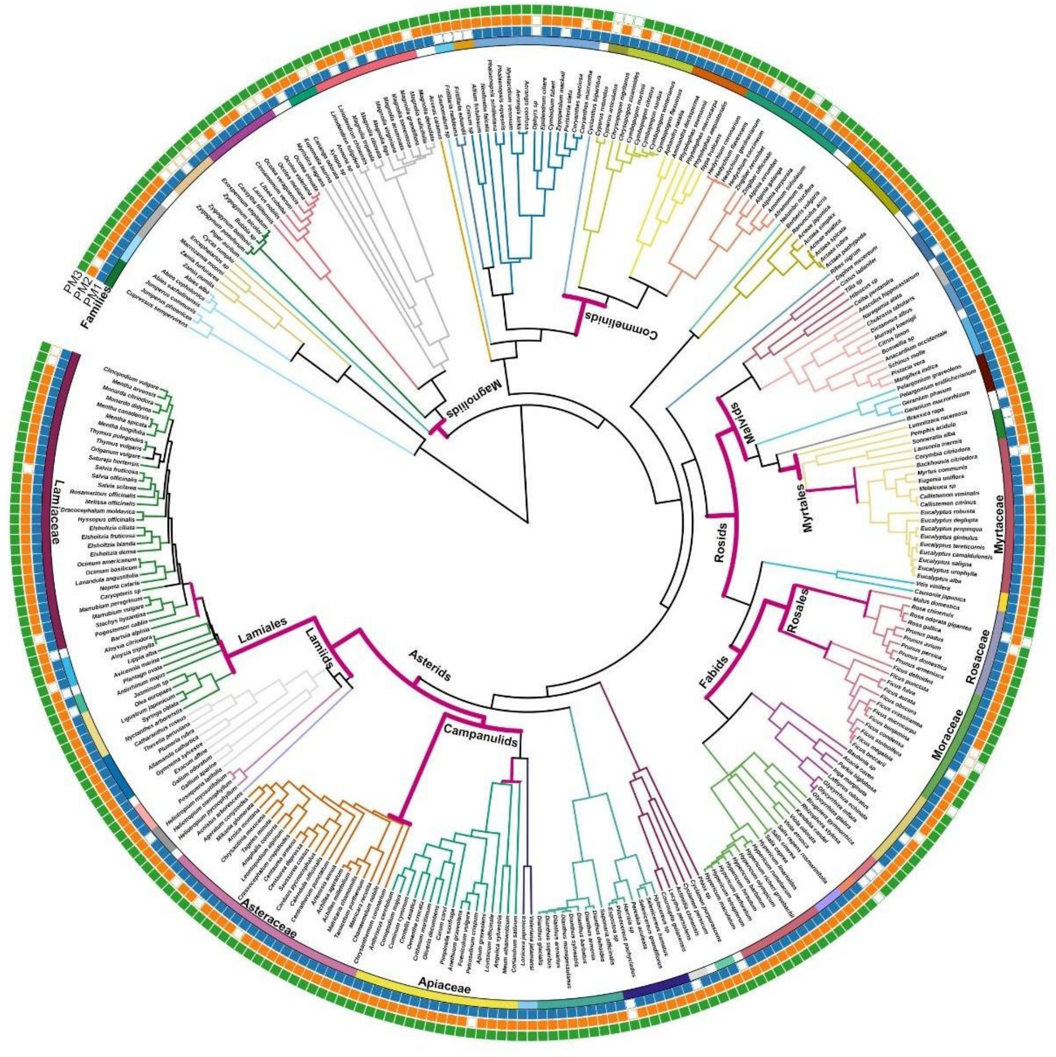


- PM1: Terpenoid dominated
- PM2: Shikimate-derived aromatics dominated
- PM3: Fatty acid derivative dominated

**Supplementary Fig. S17. Mapping of PubChem fingerprint derived clusters on to the phylogeny.** Coloured branches represent the different orders and coloured stripes represents different families. Clusters are represented as coloured squares, where presence is indicated by filled squares and absence as unfilled squares. The clades analysed for D-statistics are marked as bold magenta lines.

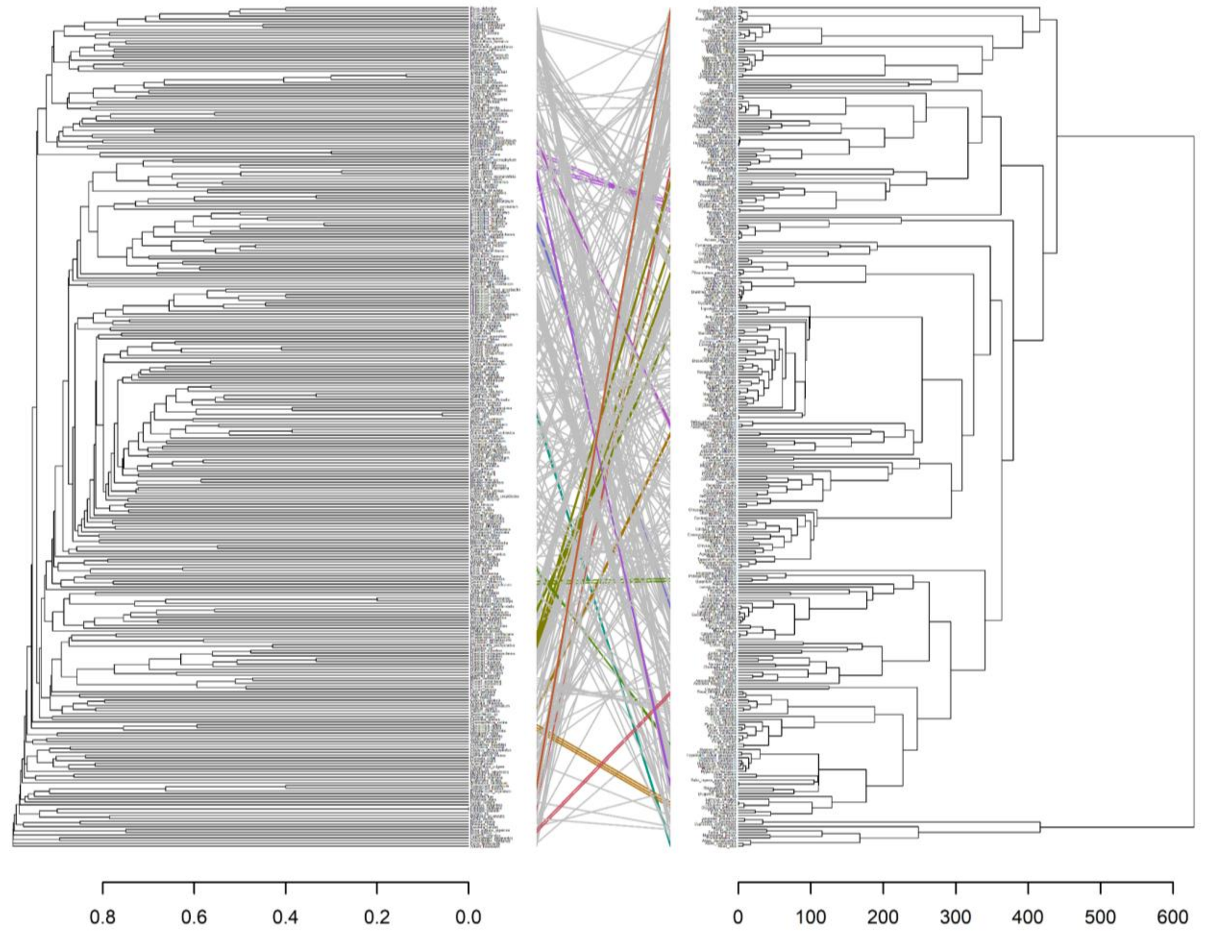


**Supplementary Fig. S18. Congruence between chemical similarity and molecular phylogeny.** Tanglegram showing the relationship between the chemical dendrogram generated from hierarchical clustering using the average linkage method (left) and molecular phylogeny using ITS sequences (right). Coloured connections represent clades that are congruent between the trees.

**Supplementary Table S1. List of abbreviations**

| | |
|---|---|
| SMILES | Simplified Molecular Input Line Entry System |
| EXT | Extended |
| MACCS | Molecular ACCess System |
| PubChem | PubChem fingerprinter |
| Sub | Substructure fingerprinter |
| SubCount | Substructure fingerprinter count |
| DBSCAN | Density-Based Spatial Clustering of Applications with Noise |
| NMI | normalised mutual information |
| ARI | adjusted rand index |
| GPP | Geranyl pyrophosphate |
| GGPP | Geranylgeranyl pyrophosphate |
| LOX | Lipoxygenase |
| MVA | Mevalonic acid |
| MEP | 2-methylerythritol 4-phosphate |
| LR | Logistic regression |
| RF | Random forest |
| MLP | MultiLayer Perceptron |
| SVC | Support Vector Classifier |
| SHAP | SHapley Additive exPlanations |
| $p_r$ | Probability of estimated D resulting from no (random) phylogenetic structure |
| $p_b$ | Probability of estimated D resulting from Brownian phylogenetic structure |
| PM | Plant metabolite cluster |
| ITS | Internal Transcribed Spacer |
| MCMC | Markov chain Monte Carlo |
| ESS | Effective sample sizes |
| MCC | Maximum clade credibility |
| ML | Machine learning |
| t-SNE | t-distributed stochastic neighbour embedding |
| GCMS | Gas chromatography- Mass spectrometry |
| GC-FID | Gas chromatography- Flame Ionisation Detector |
| iTOL | Interactive Tree Of Life |

**Supplementary Table S2.** Comparison of internal clustering validation scores obtained for different molecular fingerprint types across various clustering methods.

| | | **Silhoutte_Score** | | | | **Davies_Score** | | | | **Calinksi_Score** | | | |
|---|---|---|---|---|---|---|---|---|---|---|---|---|---|
| **Fingerprint_ Type** | **Clusters** | **kmeans** | **Agglo** | **dbScan** | **Spectral** | **kmeans** | **Agglo** | **dbScan** | **Spectral** | **kmeans** | **Agglo** | **dbScan** | **Spectral** |
| EXT | 2 | 0.15 | 0.12 | 0.2 | 0.3 | 2.3 | 2.2 | 1.9 | 2.8 | 280 | 230 | 7.4 | 13 |
| EXT | 3 | 0.15 | 0.13 | 0.31 | 0.31 | 2.2 | 2.3 | 2.2 | 2.7 | 250 | 230 | 11 | 9.6 |
| EXT | 4 | 0.14 | 0.13 | 0.32 | 0.23 | 2 | 2.1 | 2.4 | 2.9 | 220 | 210 | 15 | 7.2 |
| EXT | 5 | 0.13 | 0.13 | 0.33 | 0.28 | 2.3 | 2.2 | 2.8 | 3.3 | 190 | 180 | 21 | 9.2 |
| EXT | 6 | 0.13 | 0.1 | 0.33 | 0.25 | 2.1 | 2.2 | 2.8 | 2.8 | 170 | 160 | 21 | 6.3 |
| EXT | 7 | 0.11 | 0.11 | 0.33 | 0.23 | 2 | 2.2 | 2.8 | 2.6 | 150 | 150 | 21 | 5.1 |
| EXT | 8 | 0.13 | 0.12 | 0.33 | 0.18 | 2.1 | 2.2 | 2.8 | 1.6 | 140 | 140 | 21 | 3.9 |
| EXT | 9 | 0.15 | 0.12 | 0.33 | 0.17 | 2.1 | 2 | 2.8 | 2.2 | 140 | 130 | 21 | 5.2 |
| EXT | 10 | 0.15 | 0.12 | 0.33 | 0.21 | 2.2 | 2 | 2.8 | 3.2 | 130 | 120 | 21 | 6.3 |

| | | | | | | | | | | | | | |
|---|---|---|---|---|---|---|---|---|---|---|---|---|---|
| MACCS | 2 | 0.15 | 0.13 | 0 | 0.24 | 2.3 | 2.6 | 0 | 1.1 | 290 | 230 | 0 | 12 |
| MACCS | 3 | 0.16 | 0.13 | 0 | 0.24 | 2.1 | 2.3 | 0 | 1.2 | 270 | 210 | 0 | 13 |
| MACCS | 4 | 0.14 | 0.11 | 0 | 0.094 | 2.2 | 2.2 | 0 | 1.8 | 230 | 190 | 0 | 16 |
| MACCS | 5 | 0.14 | 0.11 | 0 | 0.077 | 2.1 | 2.1 | 0 | 1.8 | 210 | 180 | 0 | 14 |
| MACCS | 6 | 0.15 | 0.13 | 0 | 0.053 | 2.1 | 2.1 | 0 | 1.6 | 190 | 170 | 0 | 14 |
| MACCS | 7 | 0.15 | 0.13 | 0 | 0.05 | 1.9 | 2 | 0 | 1.5 | 180 | 160 | 0 | 13 |
| MACCS | 8 | 0.16 | 0.14 | 0 | 0.041 | 2 | 2.1 | 0 | 1.5 | 170 | 150 | 0 | 11 |
| MACCS | 9 | 0.17 | 0.15 | 0 | 0.011 | 2.1 | 2.1 | 0 | 1.5 | 160 | 140 | 0 | 10 |
| MACCS | 10 | 0.15 | 0.15 | 0 | 0.0064 | 2.1 | 2.1 | 0 | 1.5 | 150 | 140 | 0 | 9.3 |
| PubChem | 2 | 0.29 | 0.29 | 0 | 0.35 | 1.6 | 1.7 | 0 | 1.7 | 390 | 380 | 0 | 3.7 |
| PubChem | 3 | 0.15 | 0.14 | 0 | 0.29 | 2.1 | 2.1 | 0 | 0.53 | 330 | 310 | 0 | 3.5 |
| PubChem | 4 | 0.15 | 0.13 | 0 | 0.29 | 2 | 2 | 0 | 0.53 | 290 | 260 | 0 | 3.5 |
| PubChem | 5 | 0.16 | 0.14 | 0 | 0.29 | 2 | 1.9 | 0 | 0.53 | 260 | 240 | 0 | 3.5 |

| | | | | | | | | | | | | | |
|---|---|---|---|---|---|---|---|---|---|---|---|---|---|
| PubChem | 6 | 0.16 | 0.14 | 0 | 0.29 | 2 | 1.9 | 0 | 0.53 | 240 | 220 | 0 | 3.5 |
| PubChem | 7 | 0.16 | 0.14 | 0 | 0.29 | 2 | 2 | 0 | 0.53 | 220 | 210 | 0 | 3.5 |
| PubChem | 8 | 0.16 | 0.14 | 0 | 0.29 | 1.9 | 2 | 0 | 0.53 | 210 | 200 | 0 | 3.5 |
| PubChem | 9 | 0.16 | 0.15 | 0 | 0.29 | 1.9 | 1.9 | 0 | 0.53 | 200 | 190 | 0 | 3.5 |
| PubChem | 10 | 0.17 | 0.15 | 0 | 0.29 | 1.9 | 1.9 | 0 | 0.53 | 190 | 180 | 0 | 3.5 |
| Sub | 2 | 0.15 | 0.12 | 0 | 0.15 | 2.5 | 2.7 | 0 | 2.4 | 250 | 210 | 0 | 240 |
| Sub | 3 | 0.15 | 0.14 | 0 | 0.16 | 2.3 | 2.5 | 0 | 2 | 230 | 200 | 0 | 130 |
| Sub | 4 | 0.17 | 0.16 | 0 | 0.16 | 2 | 2.1 | 0 | 1.9 | 220 | 200 | 0 | 150 |
| Sub | 5 | 0.2 | 0.19 | 0 | 0.17 | 1.9 | 2.2 | 0 | 1.7 | 230 | 200 | 0 | 140 |
| Sub | 6 | 0.22 | 0.21 | 0 | 0.2 | 1.9 | 2 | 0 | 1.7 | 210 | 190 | 0 | 160 |
| Sub | 7 | 0.2 | 0.22 | 0 | 0.21 | 2 | 1.9 | 0 | 1.6 | 190 | 180 | 0 | 150 |
| Sub | 8 | 0.21 | 0.22 | 0 | 0.21 | 1.9 | 1.9 | 0 | 1.6 | 190 | 170 | 0 | 140 |
| Sub | 9 | 0.22 | 0.19 | 0 | 0.21 | 1.8 | 1.9 | 0 | 1.6 | 180 | 160 | 0 | 130 |

| | | | | | | | | | | | | | |
|---|---|---|---|---|---|---|---|---|---|---|---|---|---|
| Sub | 10 | 0.25 | 0.2 | 0 | 0.23 | 1.7 | 1.9 | 0 | 1.6 | 170 | 160 | 0 | 130 |
| Sub_count | 2 | 0.37 | 0.37 | 0.58 | 0.36 | 1.1 | 1.1 | 1.2 | 0.5 | 1100 | 1100 | 16 | 2.9 |
| Sub_count | 3 | 0.36 | 0.35 | 0.58 | 0.29 | 0.94 | 0.89 | 1.2 | 0.88 | 1300 | 1100 | 16 | 3.7 |
| Sub_count | 4 | 0.34 | 0.32 | 0.58 | 0.32 | 0.99 | 1 | 1.2 | 1.3 | 1300 | 1100 | 16 | 10 |
| Sub_count | 5 | 0.36 | 0.34 | 0.58 | 0.36 | 0.91 | 0.92 | 1.2 | 3.3 | 1300 | 1200 | 16 | 5.9 |
| Sub_count | 6 | 0.31 | 0.28 | 0.58 | 0.27 | 1 | 1 | 1.2 | 2 | 1200 | 1100 | 16 | 5.4 |
| Sub_count | 7 | 0.32 | 0.26 | 0.58 | 0.31 | 1 | 1.2 | 1.2 | 0.41 | 1200 | 1000 | 16 | 9.6 |
| Sub_count | 8 | 0.29 | 0.26 | 0.58 | 0.28 | 1.1 | 1.2 | 1.2 | 0.93 | 1100 | 970 | 16 | 8 |
| Sub_count | 9 | 0.28 | 0.23 | 0.58 | 0.23 | 1.2 | 1.2 | 1.2 | 0.52 | 1100 | 940 | 16 | 2.9 |
| Sub_count | 10 | 0.26 | 0.24 | 0.58 | 0.34 | 1.2 | 1.2 | 1.2 | 1.6 | 1000 | 910 | 16 | 6.2 |

**Supplementary Table S3. Phylogenetic signals calculated as D-statistics for SubCount and PubChem fingerprints across seed plants and selected angiosperm clades.** A value of D = 0 means phylogenetically conserved as expected under Brownian threshold motion, while D = 1 indicates random distribution of the trait across the phylogeny. Values of D < 0 indicate stronger phylogenetic clustering (traits more conserved than expected under Brownian motion), while values of D > 1 mean phylogenetic overdispersion (traits more divergent than random). $p < 0.05$ means the D-value is statistically significant and is represented in bold. $p_r$ = Probability of estimated D resulting from no (random) phylogenetic structure; $p_b$ = Probability of estimated D resulting from Brownian phylogenetic structure, PM = Plant metabolite cluster.

| FP type | Clusters | Whole tree (D-statistic $p_r/p_b$) | Rosids (D-statistic $p_r/p_b$) | Asterids (D-statistic $p_r/p_b$) | Malvids (D-statistic $p_r/p_b$) | Fabids (D-statistic $p_r/p_b$) | Lamids (D-statistic $p_r/p_b$) | Campanulids (D-statistic $p_r/p_b$) |
|---|---|---|---|---|---|---|---|---|
| SubCount | PM1a: Sesqui- and diterpene dominated | 0.6112 (**0**/**0**) | 0.8444 (0.153/**0**) | 0.4440 (**0.002**/0.084) | 0.7010 (0.158/0.097) | 0.8436 (0.211/**0.012**) | -0.1812 (**0**/0.664) | 1.1614 (0.614/**0.017**) |
| | PM1b: Monoterpene dominated | 0.6578 (**0**/**0**) | 0.4364 (**0**/**0.046**) | 0.3463 (**0.001**/0.165) | 0.2384 (**0.029**/0.37) | 0.4375 (**0.004**/0.089) | 0.6549 (0.097/0.093) | NA |
| | PM2: Simple benzenoids and phenylpropanoids + Aromatic nitrogen and sulphur containing compounds dominated | 0.7595 (**0.001**/**0**) | 0.8190 (0.158/**0.007**) | 0.9264 (0.308/**0**) | 1.4425 (0.92/**0.001**) | 0.1724 (**0.011**/0.373) | 0.8459 (0.292/0.082) | 0.9800 (0.456/**0.024**) |
| | PM3a: Aliphatics and C5 branched chain compounds dominated | 0.6566 (**0**/**0**) | 0.4524 (**0**/**0.015**) | 0.7761 (**0.044**/**0**) | 0.8195 (0.244/**0.019**) | 0.2167 (**0**/0.219) | 0.4899 (**0.018**/0.082) | 0.9926 (0.464/**0.001**) |
| | PM3b: Aliphatics and C5 branched chain compounds dominated | 0.3823 (**0**/**0.007**) | -0.0815 (**0**/0.619) | 0.5643 (**0.002**/**0.027**) | 0.3069 (**0.027**/0.31) | -0.4620 (**0**/0.914) | 0.3143 (**0.005**/0.267) | 1.2879 (0.722/**0.016**) |

| | | | | | | | | |
|---|---|---|---|---|---|---|---|---|
| | PM3c: Non-aromatic nitrogen and sulphur containing compounds + Aliphatics and C5 branched chain compounds dominated | 0.6802 (**0**/**0**) | 0.5269 (**0**/**0.005**) | 0.7044 (**0.024**/**0.003**) | 0.9948 (0.46/**0.003**) | -0.0875 (**0**/0.598) | 0.5148 (**0.03**/0.097) | 0.6527 (0.06/0.073) |
| | PM3d: Aliphatics and C5 branched chain compounds dominated | 1.0166 (0.542/**0**) | 1.2106 (0.857/**0**) | 1.0915 (0.642/**0.001**) | 0.6739 (0.225/0.255) | 1.4281 (0.973/**0**) | 1.1959 (0.68/**0.013**) | 1.1513 (0.626/**0.011**) |
| PubChem | PM1: Terpenoid dominated | 0.6709 (**0**/**0**) | 0.6251 (**0.033**/**0.034**) | 0.3367 (**0.013**/0.29) | 0.6319 (0.145/0.149) | 0.3466 (**0.019**/0.239) | 0.2292 (**0.042**/0.395) | NA |
| | PM2: Shikimate-derived aromatics dominated | 0.6861 (**0**/**0**) | 0.4151 (**0.013**/0.191) | 1.0236 (0.513/**0.003**) | 0.8899 (0.389/0.247) | 0.1724 (**0.011**/0.373) | 0.8459 (0.292/0.082) | 1.2948 (0.712/**0.015**) |
| | PM3: Fatty acid derivative dominated | 0.571 (**0.001**/0.061) | 1.2436 (0.827/**0.003**) | NA | NA | 1.4498 (0.952/**0**) | NA | NA |

**Continued**

| FP type | Clusters | Commelinids (D-statistic $p_r/p_b$) | Magnoliids (D-statistic $p_r/p_b$) | Myrtales (D-statistic $p_r/p_b$) | Rosales (D-statistic $p_r/p_b$) | Lamiales (D-statistic $p_r/p_b$) | Asteraceae (D-statistic $p_r/p_b$) |
|---|---|---|---|---|---|---|---|
| SubCount | PM1a: Sesqui- and diterpene dominated | 1.3063 (0.677/0.069) | 0.4279 (**0.012**/0.109) | 0.5445 (0.174/0.245) | 1.0189 (0.484/**0.024**) | -0.2366 (**0.012**/0.647) | 2.069 (0.514/0.285) |
| | PM1b: Monoterpene dominated | 0.4895 (0.085/0.188) | 0.5972 (0.076/0.079) | -0.1709 (**0.014**/0.624) | 0.6468 (0.146/0.107) | 0.3155 (0.054/0.34) | NA |

| | | | | | | | |
|---|---|---|---|---|---|---|---|
| | PM2: Simple benzenoids and phenylpropanoids + Aromatic nitrogen and sulphur containing compounds dominated | 0.6241 (0.128/0.1) | 0.7195 (0.113/**0.018**) | 1.0843 (0.421/0.202) | -0.2017 (**0.013**/0.627) | 0.9352 (0.371/0.063) | 1.2800 (0.69/**0.022**) |
| | PM3a: Aliphatics and C5 branched chain compounds dominated | 0.5336 (0.061/0.141) | 0.5756 (**0.043**/0.064) | 0.7337 (0.289/0.214) | 0.2528 (**0.021**/0.305) | 0.5479 (0.085/**0.009**) | 0.4398 (0.129/0.254) |
| | PM3b: Aliphatics and C5 branched chain compounds dominated | 4.9933 (0.737/0.177) | NA | NA | NA | 0.8682 (0.321/0.095) | 1.0168 (0.423/0.246) |
| | PM3c: Non-aromatic nitrogen and sulphur containing compounds + Aliphatics and C5 branched chain compounds dominated | 1.0972 (0.532/**0.035**) | 0.6959 (0.094/**0.024**) | 1.020 (0.456/**0.045**) | -0.7788 (**0**/0.955) | 0.5434 (0.087/0.123) | 0.6938 (0.248/0.152) |
| | PM3d: Aliphatics and C5 branched chain compounds dominated | 0.9349 (0.387/**0.015**) | 0.6254 (0.172/0.15) | NA | 1.8404 (0.988/**0**) | 1.0012 (0.433/0.112) | 1.0274 (0.457/0.137) |
| PubChem | PM1: Terpenoid dominated | NA | 0.7371 (0.197/0.078) | 0.3011 (0.114/0.411) | 1.6687 (0.812/**0.024**) | -0.4698 (**0.018**/0.73) | NA |
| | PM2: Shikimate-derived aromatics dominated | 0.6241 (0.128/0.1) | 0.7195 (0.113/**0.018**) | 1.0843 (0.421/0.202) | -0.2017 (**0.013**/0.627) | 0.9352 (0.371/0.063) | 1.9097 (0.886/**0.009**) |
| | PM3: Fatty acid derivative dominated | 0.1782 (**0.012**/0.372) | NA | NA | 1.8404 (0.988/**0**) | NA | NA |

**Continued**

| FP type | Clusters | Lamiaceae (D-statistic $p_r/p_b$) | Moraceae (D-statistic $p_r/p_b$) | Rosaceae (D-statistic $p_r/p_b$) | Myrtaceae (D-statistic $p_r/p_b$) | Apiaceae (D-statistic $p_r/p_b$) |
|---|---|---|---|---|---|---|
| SubCount | PM1a: Sesqui- and diterpene dominated | NA | 1.981 (0.924/**0.007**) | -2.0107 (0.02/0.855) | 0.1332 (0.119/0.474) | 1.6189 (0.757/**0.039**) |
| | PM1b: Monoterpene dominated | -0.8690 (0.025/0.738) | 0.6526 (0.292/0.214) | -2.0107 (0.02/0.855) | -0.6969 (0.067/0.71) | NA |
| | PM2: Simple benzenoids and phenylpropanoids + Aromatic nitrogen and sulphur containing compounds dominated | 0.5488 (0.148/0.282) | -1.8033 (0/0.957) | 2.2798 (0.543/0.191) | 0.8212 (0.354/0.233) | -0.4532 (0.185/0.71) |
| | PM3a: Aliphatics and C5 branched chain compounds dominated | 0.5146 (0.124/0.173) | 2.135 (0.843/**0**) | 0.3779 (0.194/0.413) | 1.5576 (0.624/0.294) | 1.432 (0.717/**0.023**) |
| | PM3b: Aliphatics and C5 branched chain compounds dominated | 1.0819 (0.44/0.24) | NA | NA | NA | 2.777 (0.886/0.084) |
| | PM3c: Non-aromatic nitrogen and sulphur containing compounds + Aliphatics and C5 branched chain compounds dominated | 0.6950 (0.222/0.159) | NA | -2.0107 (0.02/0.855) | 0.8318 (0.352/0.129) | 0.0615 (0.086/0.545) |

| | | | | | | |
|---|---|---|---|---|---|---|
| | PM3d: Aliphatics and C5 branched chain compounds dominated | 1.3035 (0.569/0.18) | 2.3109 (0.955/**0**) | 2.0477 (0.517/0.191) | NA | 1.7154 (0.694/0.136) |
| PubChem | PM1: Terpenoid dominated | NA | 1.5517 (0.51/0.189) | 2.0477 (0.517/0.191) | -0.6969 (0.067/0.71) | NA |
| | PM2: Shikimate-derived aromatics dominated | 0.5488 (0.148/0.282) | -1.8033 (0/0.957) | 2.2798 (0.543/0.191) | 0.8212 (0.354/0.233) | -0.4532 (0.185/0.71) |
| | PM3: Fatty acid derivative dominated | NA | 2.3109 (0.955/**0**) | 2.0477 (0.517/0.191) | NA | NA |

**Supplementary Table S4. Feature filtering methods**

| Feature Set | Before filtering | After Jaccard similarity- based selection | After correlation-based selection | After filtering near-constant features |
|---|---|---|---|---|
| Volatile metabolites | 2139 | 2139 | 1215 | 401 |
| Volatile metabolites + EXT | 3163 | 3163 | 2007 | 522 |
| Volatile metabolites + MACCS | 2305 | 2305 | 1348 | 441 |
| Volatile metabolites + PubChem | 3020 | 3020 | 1841 | 455 |
| Volatile metabolites + Sub Count | 2446 | 2446 | 1495 | 419 |
| Volatile metabolites + Sub | 2446 | 2446 | 1495 | 415 |

**Supplementary Note S1. Feature filtering protocol**

First, we applied Jaccard similarity-based selection, which helps identify and remove redundant or highly similar features by measuring the overlap of feature sets. However, no features were eliminated for the considered threshold of 0.9. Next, we applied a correlation-based selection method, which helps eliminate strongly correlated features, thereby reducing multicollinearity and improving the stability of the model. For this, we used a threshold of 0.9, and any feature that had a correlation higher than this was removed. Finally, we applied filtering of near-constant features, that is, those with little to no variability across samples and that typically contribute little to the predictive performance. In this case, we used a variance threshold of 0.01, and features with less variance were removed. This filtering contributes the most to feature reduction in all datasets.

**Supplementary Note S2. Detailed description of classification models**

LR is a traditional statistical technique that fits a straight line to learn the relationship between input and output variables. In contrast, RF is an ensemble method that constructs numerous decision trees and aggregates their results to increase accuracy and robustness, particularly in cases where the data contain intricate or erratic patterns. SVC is based on Support Vector Machines (SVMs), which operate by determining the best hyperplane with the largest margin between data points of various classifications. By using kernel functions (such as linear, polynomial, and radial basis functions), which convert the data into higher-dimensional spaces for improved separation, SVC can handle both linear and non-linear decision boundaries. It is effective in high-dimensional settings, robust to overfitting when properly regularised, and particularly useful when the number of features is large compared to the number of samples. MLP is a type of artificial neural network that consists of layers of interconnected nodes and is capable of capturing more intricate, non-linear relationships, making it a versatile tool for classification tasks.

**Supplementary Note S3. Evaluation of confusion matrices from classification models**

The confusion matrix identifies samples that are correctly predicted to true classes or misclassified into different classes. Higher values in the diagonals of the matrix (marked as red boxes) represent a perfect match, whereas zeroes in the diagonal represent misclassifications. The misclassifications might arise due to no unique chemical signal detectable by the model, overlapping chemistries with other classes or too less samples to interpret. If the overlapping chemistry reflects closely related groups, that suggests a phylogenetic constraint, while if the groups are unrelated and still show chemical similarity, given some ecological or environmental similarity, it might indicate convergent evolution.

**Supplementary Note S4. Important features that helped in the classification models**

The top 400 features, or volatile metabolites, based on the importance scores, were evaluated for the best-performing model per rank and were examined to determine if they represented the metabolites that were unique to each of the taxonomic ranks. Most of the features are non-unique. While a very few unique compounds exist, their importance values are zero or less than 0.01.

**Supplementary Note S5. Feature importance using SHAP (SHapley Additive exPlanations) heatmaps**

The SHAP heatmaps provide insights into how different features contribute to predictions across multiple ranks. Each row represents a feature, and each column corresponds to an individual instance or a datapoint. The colour intensity (red vs. blue) indicates the magnitude and direction of feature influence: red suggests a feature is pushing the prediction toward a certain taxonomic rank, while blue indicates it is pulling the prediction away from that rank. The addition of molecular fingerprints consistently simplifies the learning, allowing the models to find a "cleaner" and more focused path to the answer, as seen in the order- and family-level classification. Without the fingerprints, the models rely on "busier" and more complex logic, weighing a much broader combination of compounds to find a pattern. This feature dependency also dictates which compounds the models find important. Two different feature sets caused the models to identify completely different chemicals as the key positive signals.

**Supplementary Note S6. Tree-building protocols**

**Molecular markers used for the phylogenetic reconstruction:**

Internal Transcribed Spacer (ITS) sequences for 305 available plant species were downloaded from the NCBI Genbank (https://www.ncbi.nlm.nih.gov/nucleotide/). The sequences were aligned using ClustalW in Geneious 10.0.1. A maximum likelihood tree was generated using IQ Tree on the CIPRES portal (https://www.phylo.org/). Genetic distances were extracted with the cophenetic.phylo function ("ape" package).

A time-calibrated Bayesian phylogeny was then estimated in BEAST 2.7.7 using the GTR+I+J model (determined from the JModel test), an uncorrelated log-normal relaxed clock, and 33 fossil calibration points. We executed five independent Markov chain Monte Carlo (MCMC) runs, each comprising 100 million generations with sampling every 100 generations. We combined the five independent runs using the LogCombiner[3], with a burn-in set to 10% of the initial samples from each run, resampling at lower frequencies (5000 trees) and verifying the effective sample sizes (ESS > 200) in Tracer version 1.7.2[4]. Further, we employed the TreeAnnotator[5] to derive the maximum clade credibility (MCC) tree.

**Supplementary Note S7. Estimation of the chemical versus genetic congruence**

The binary volatile metabolite dataset was aggregated at the species level, with presence recorded if the compound appeared at least once in any sample of the same species. Species-level chemical dissimilarities were calculated as pairwise Jaccard distance using the vegdist function ("vegan" package). Species-level genetic dissimilarity was estimated from the maximum likelihood tree using the cophenetic.phylo function ("ape" package). Furthermore, the Spearman correlation between the chemical and genetic distances was assessed through Mantel's test, implemented using the mantel function with 999 permutations.

To evaluate topological similarities, a chemical dendrogram generated via hierarchical clustering (average linkage method) using the hclust function and the dated MCC tree (pruned to common tips) was compared using Baker's gamma correlation coefficient using the cor_bakers_gamma function in the "dendextend" package. All the analyses were performed in R version 4.5.1[6].

**Results:**

The Mantel's test revealed a weak but statistically significant positive correlation between the chemical distance at the species level and genetic distance from the maximum likelihood tree (Mantel's $r = 0.0898$, $p = 0.007$). The chemical constraints in some of the sister lineages were visualised via topological congruences between the chemical dendrogram based on the average linkage method and Bayesian phylogeny (Supplementary Fig. 16). This was also supported by the very weak yet significant Baker's gamma correlation coefficient (Baker's gamma = 0.0966, $p = 0.01$ based on 99 permutations). The observed correlation is therefore unlikely to have occurred by chance, thus indicating a weak phylogenetic signal in the composition of volatile metabolites of the studied plant species.
